# Droplet Sequencing Reveals Virulence Gene Clusters in Oral Biofilm Extracellular Vesicles

**DOI:** 10.1101/2024.09.18.613607

**Authors:** Sotaro Takano, Naradasu Divya, Satoshi Takenawa, Yan Kangmin, Tomoko Maehara, Nobuhiko Nomura, Nozomu Obana, Masanori Toyofuku, Michihiko Usui, Wataru Ariyoshi, Akihiro Okamoto

## Abstract

Bacterial extracellular vesicles (BEVs), produced by a broad spectrum of bacteria, play a crucial role in enhancing intercellular communication through DNA transfer. A vital determinant of their gene transfer efficiency is the gene content diversity within BEVs, an aspect that conventional metagenomics fails to capture. Our study bridges this gap with a novel microdroplet-based sequencing technique that precisely details DNA content within hundreds of individual BEVs. This technique revealed a unique DNA profile in BEVs from the oral pathogen *Porphyromonas gingivalis*, pinpointing specific genomic regions related to DNA integration (e.g., DNA transposition and CRISPR-Cas systems). These enriched genes, overlooked by standard analyses that aggregate total read counts, indicate that our method offers a more focused view into the genetic contents of BEVs. Applying our technique to dental plaque-derived BEVs, we discovered a hundredfold higher prevalence of DNA encapsulation than previously estimated, with over 30% of BEVs containing DNA. Specifically, we identified a substantial presence of O-antigen biosynthesis genes, prominent hotspots of frequent horizontal gene transfer, from *Alcaligenes faecalis*. Given that O-antigens mediate host-bacterial interactions, this gene enrichment in the large fraction of BEVs suggests a potential novel pathway by which BEVs could influence pathogenicity within oral biofilms. Our research unveils critical insights into the potential functions of vesicular DNA in microbial communities, establishing a powerful platform for studying vesicular DNA in microbiomes. This technical breakthrough provides a foundational basis for future research in microbial communication and the development of potential therapeutic or diagnosis strategies.

**Significance Statement:** BEVs have been studied for decades, yet their roles in nature and disease are just beginning to be appreciated. Our study makes a significant leap in understanding the roles of BEVs as gene transfer vehicles. By developing a microdroplet-based sequencing technique, we have uncovered detailed DNA profiles within individual BEVs, a task beyond the capabilities of conventional metagenomic methods. This breakthrough highlights specific genomic regions enriched in BEVs from pure culture and human dental plaque. Furthermore, the high prevalence of biofilm BEVs enriched in O-antigen biosynthesis genes, suggests a potential impact on the pathogenicity of oral biofilms. This research establishes a new methodological platform for exploring the intricacies of BEV-mediated interactions in a complex microbial community.

## Introduction

Horizontal gene transfer (HGT), the exchange of genetic materials across different organisms, represents a fundamental mechanism in microbial evolution (1). This process enables bacterial cells to acquire novel phenotypic traits (e.g., antibiotic resistance), fostering rapid adaptations to environmental challenges (2–4). HGT’s role extends beyond individual survival, significantly influencing the genetic diversification within microbial populations (5). It allows the sharing and integration of beneficial genes across species barriers (6, 7), catalyzing the emergence of novel capabilities within bacterial communities (2, 8). Bacterial extracellular vesicles (BEVs) have emerged as critical facilitators of HGT (9, 10). These vesicles are produced by both Gram-positive and Gram-negative bacteria, encapsulate and condense diverse biomolecules including DNA (11), and are prevalent in the various habitats (12, 13). The internal molecules in BEVs are generally condensed, secreted and protected from the risks of degradation (14), hence the success rate of gene transfer increases in BEV-mediated transfer (15). BEVs have the potential to transfer genetic material not only within a species but also across phylogenetically distant species (16–18). These characteristics are either superior or complement to other molecular mechanisms (e.g., conjugation and natural transformation), expanding the genetic repertoire and adaptability in microbial communities. However, the specific nature of the genetic material within BEVs has remained largely underexplored, obscured by technical limitations in current genomic analysis methods.

A critical factor determining the effectiveness of HGT is the quantity of BEVs containing specific genes (16), underscoring the need to understand which genes are present and the diversity of DNA encapsulated in individual BEVs. However, identifying and characterizing the DNA within BEVs poses significant challenges, as conventional metagenomic methods do not adequately differentiate DNA variations among individual vesicles. BEVs selectively enrich certain DNA pieces (19–21) but vesicles from the same bacterial culture can contain different DNA (22). To address this, we apply a novel single-nanoparticle droplet DNA sequencing (NP-DS) technique, adapted from single-cell genomic methodologies (23–25). Applied to BEVs from *Porphyromonas gingivalis* and human dental plaque, genetic contents in individual vesicles were profiled. Given this organism and its BEVs have been implicated in periodontitis and systemic diseases (26, 27), our study points to specific aspects of BEV-derived DNA that may play important functional roles in microbial interactions, particularly in the context of oral biofilms and infection.

## Results

### Enrichment of specific genomic regions in BEVs produced by an oral pathogenic bacterium

We first explored the characteristics of DNA sequences in BEVs produced by *P. gingivalis,* a prominent pathogen of periodontitis (28, 29). Nanoparticle tracking analysis (NTA) of the isolated samples detected > 10^9^ particles (/mL) stained by lipophilic dye (Fig. S1A), indicating the presence of BEVs in bacterial cell cultures. To understand the composition of the DNA housed within these vesicles, we performed shotgun sequencing for the bulk BEV samples (Fig. 1A), after treating the BEVs with DNase I to remove any external DNA. The sequence reads were widely mapped to the original bacterial genome, but several genomic regions were over-detected (Fig. 1B), suggesting a selective DNA packaging within the BEV population.

**Figure 1.**
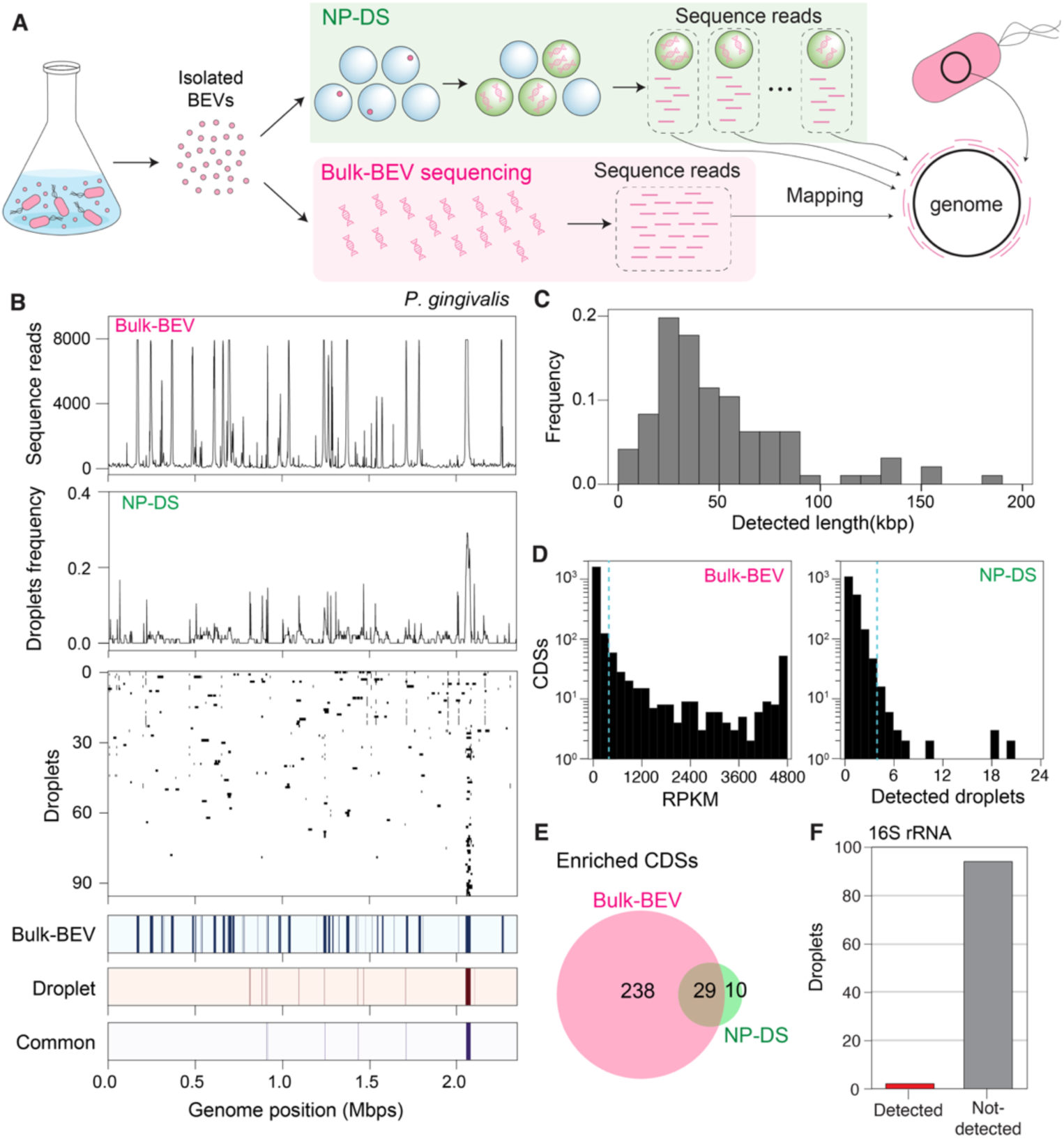
Enrichment of DNA at specific loci of the host genome in *P. gingivalis* BEVs. (A) A schematic of the analysis. For the isolated BEVs from the bacterial culture, we performed nanoparticle droplet sequencing (NP-DS) by DNA amplification of encapsulated droplets or bulk-BEV sequencing by directly extracting DNA from the whole BEV particles for shotgun sequencing. In NP-DS, the collected sequence reads in each droplet was mapped to the original bacterial genome to obtain mapping profiles of individual droplets. In bulk-BEV sequencing, all sequence reads were mapped to the genome collectively. (B) Loci of mapped regions by sequence reads in BEVs on the originating bacterial chromosomes. The number of mapped reads in the next-generation sequencing of the bulk-BEV sample was displayed in a top panel. In the second top panel, the frequency of positive droplets in 96 across each bacterial genome is shown. In the third top panel, the positions on the genome where the sequence reads were mapped in each droplet were filled with black. The genomic regions determined as significantly enriched in bulk-BEVs (more than 75 percentile + 2*IQR (interquartile range)) or droplet sequencing (binominal test, p < 0.01) were filled with red and blue in the bottom panels, respectively. The enriched chromosomal regions in both the droplet and bulk-BEVs sequence analysis were filled with purple (in the bottom-most panel). (C) The distributions of total length of detected CDSs. (D) The abundance distribution of 1873 CDSs in BEVs. In the bulk-BEV sequencing, the abundance was estimated by the frequency of mapped reads (RPKM). In the case of the NP-DS, the number of droplets detected was used as a metric of the abundance of each CDS. Blue dashed lines indicate the threshold for enriched CDSs (i.e., outliers, 75 percentile + 2*IQR (interquartile range)). (E) Venn diagram for the number of enriched CDSs in the bulk-BEV sequencing and the NP-DS. (F) The percentage of BEV-containing droplets in which 16S rRNA sequence was detected and not detected. The DNA sample is the mixture of BEVs collected from three independent cultures.

To test this possibility, we performed whole genome amplification (WGA) for those particles by droplet-based method (23). Each nanoparticle was isolated within a gel bead, facilitating in-gel lysis and WGA (see Materials and Methods). The sample was mixed with agarose so that ≈ 40 % of droplets contained a single nanoparticle (Table 1, Fig. S2, presented as *R_t_*). This process harbored 4.6 % of the droplets contained amplified DNA (stained by SYBR-Green-I, Table 1, Fig. S2, presented as *Ro*). Given that nearly 20 % of the total nanoparticles were lipid-stained (Table 1, Fig. S1 and Fig S2, presented as *R_b_*), we could estimate that ≈ 60 % of lipid-stained nanoparticles (BEVs) harbored DNA (Table 1). This substantial population underscores a prevalent DNA packaging process within the BEVs of *P. gingivalis*, highly surpassing the low DNA packaging frequency in BEVs (<1%) previously estimated (22). Using SYBR-Green-I (the dye for nucleic acid) directly to BEVs as a conventional method, we detected only less than 1% of positively stained BEVs by NTA (Fig. S1), which is consistent with previous pure culture study (22). Considering that fluorescence staining sensitivity is contingent on DNA length (30), it was conceivable that most of BEVs might contain DNA segments too short to be detected by this technique, and there would be considerable variations in DNA length among BEV particles.

**Table 1.**
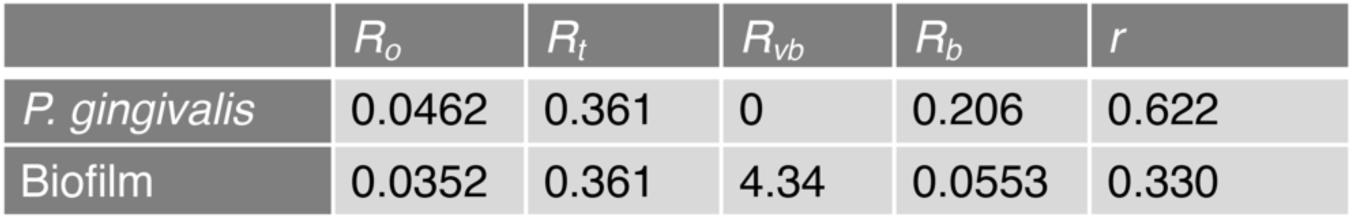
The percentage of droplets containing amplified DNA fragments after processing cells or nanoparticles. The ratio of droplets to particles with DNA (*R_o_*). The theoretical ratio of droplets to total particles (*R_t_*) was controlled to a fixed value in our analysis. *R_o_* and *R_vb_* are estimated by NTA and taxonomic annotation, we calculated the percentage of BEVs with DNA, *r*. For calculation details, see Supporting Materials and Methods and Figure S2.

| | $R_o$ | $R_t$ | $R_{vb}$ | $R_b$ | $r$ |
| --- | --- | --- | --- | --- | --- |
| *P. gingivalis* | 0.0462 | 0.361 | 0 | 0.206 | 0.622 |
| Biofilm | 0.0352 | 0.361 | 4.34 | 0.0553 | 0.330 |

To elucidate the DNA content within individual BEV particles, we performed whole-genome sequencing for the DNA amplified by multiple displacement analysis (MDA). We examined the distribution of sequence reads from 96 droplets and found that wide regions of the host chromosome were detected in at least one of the droplets same as bulk-BEV sequencing (Fig. 1B). However, there were variations in the detected genomic regions among BEV-population in *P. gingivalis* (Fig. 1B) and the averagely 51 ± 35 kbp of genomic regions were detected in each droplet (Fig. 1C and Table. S1).

While the mapped regions by read sequences from *P. gingivalis*-BEVs were not uniformly distributed among 96 droplets (Fig. 1B), we found a few genomic regions that were significantly over-detected among BEV population (binominal test, p < 0.01, see Supporting Materials and Methods) (Fig. 1B). Although there were commonly enriched regions between the bulk-BEV-sequencing and the droplet sequencing of BEVs in *P. gingivalis* (Fig. 1B), some of the frequently mapped regions in bulk-BEV sequencing were not classified as enriched in the criterion of the detected droplets. The deviation between the two approaches was also observed in the prevalence of each protein coding sequences (CDSs) among BEVs. The abundance distribution for 1873 CDSs estimated by mapped sequence reads (FPKM) in the bulk-BEV-sequencing is longer-tailed than that estimated by the detected droplets in the NP-DS (Fig. 1D). Consequently, the bulk-BEV-sequencing harbored ≈ 7 times more CDSs regarded as enriched (i.e., outliers in the distribution, see Data S1) than the NP-DS (Fig. 1E). Additionally, 16S rRNA sequence was rarely detected in BEVs of *P. gingivalis* (≈ 2 %, Fig. 1F), suggesting that detecting BEVs by this marker gene is less efficient compared to the other enriched genomic regions in this bacterium. Of note, MDA usually harbors chimeric sequences during the amplification process (31), but we confirmed that such chimeric reads did not harbor qualitative difference in the mapping profiles (Fig. S3 and see Supporting Materials and Methods for details).

### Enrichment of functional genes in BEVs from oral pathogen

We next probed the functional implications of DNA enrichment within *Porphyromonas gingivalis* BEVs. To explore the genes loaded into a large fraction of BEV population, we statistically extracted the over-detected CDSs among 96 droplets in the NP-DS, revealing 39 significantly over-represented CDSs out of the 1873 present on the genome (Table S1). Around 20 % of the enriched CDSs are related to genome arrangements such as transposase, site-specific integrase, and CRISPR-Cas system (9/39).

To further dissect their function, we performed homology search of those enriched CDSs against the UniprotKB/Swiss-Prot database (32), categorized them by Gene Ontology terms (GO terms), and statistically screened the enriched GO terms in BEVs (Fig. 2A and Table S2). Among 497 categories, 5 categories were identified as statistically enriched (hypergeometric test, p-value < 0.05, see Supporting Materials and Methods). We found that “maintenance of CRISPR repeat elements (GO:0004803)” and “DNA transposition (GO:0006313)” emerged as statistically significant categories, indicating the predominance of genome-arrangement functions in the BEV-derived DNA. The “maintenance of CRISPR repeat elements” category, which is also linked to the “defense response to virus (GO:0051607)”, was particularly highlighted by the packaging of an entire CRISPR-Cas gene cluster within an enriched genomic region (Fig. 2B). Also, “cobalamin biosynthetic process (GO: 0009236)”, was identified as the enriched functional category (Fig. 2A). All four genes associated with this metabolism are not distributed across the genome but are clustered on the specific locus (Fig. 2C).

**Figure 2.**
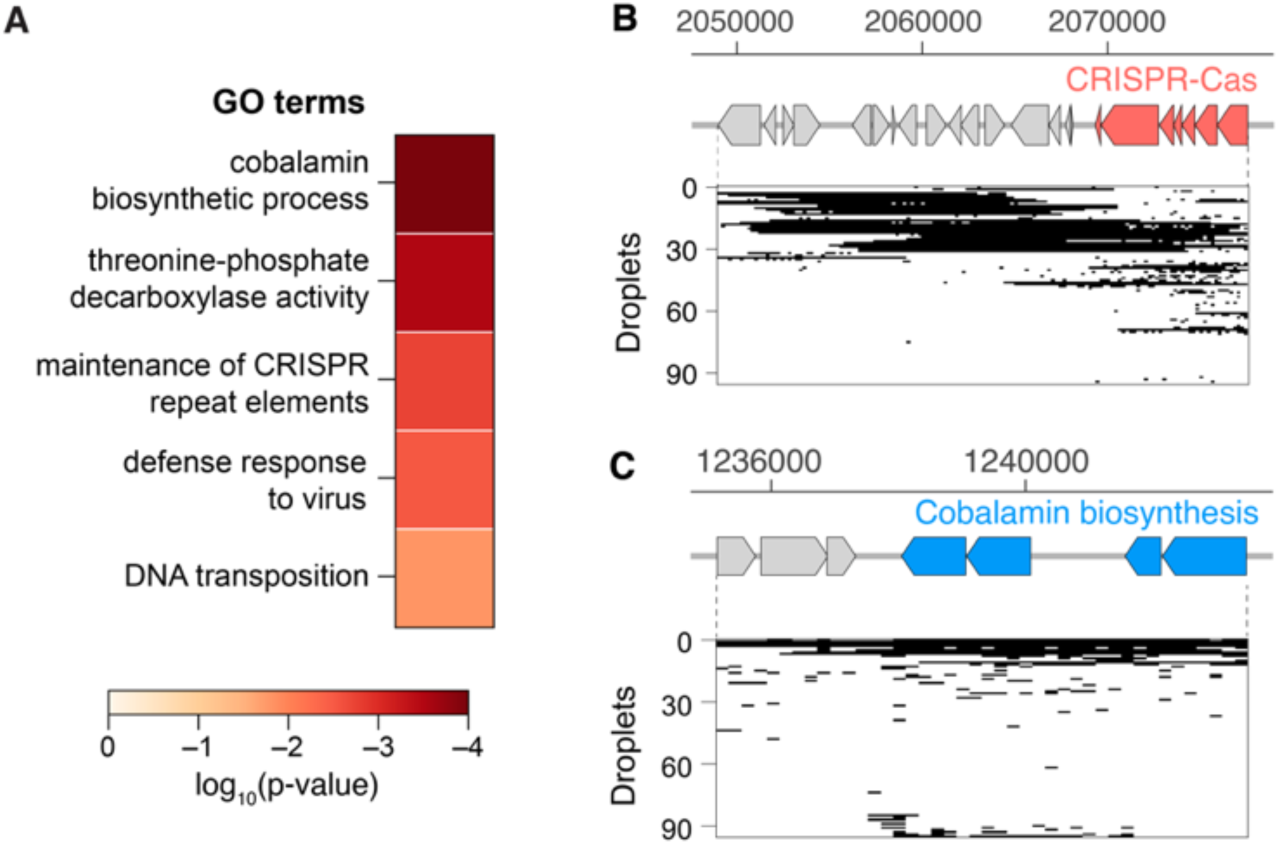
Functional enrichment of genes in *P. gingivalis* BEVs. (A) Enriched gene categories in CDSs detected in BEVs of *P. gingivalis*. Frequently detected CDSs in BEVs were statistically extracted (hypergeometric test, p-value < 0.05, see Supporting Materials and Methods). Gene ontology terms (GO terms) were used for the classification of those frequently detected CDSs. Heatmap indicates the statistical significance level of enrichment. (B), (C) Clustered functional genes located on the enriched genomic regions in *P. gingivalis* BEVs. Detection profiles in 96 droplets where (B) CRISPR-Cas gene cluster or (C) cobalamin biosynthetic genes located.

We also performed the same analysis of genes in bulk-BEV sequencing and confirmed that the same or similar GO terms such as “cobalamin biosynthetic process (GO: 0009236)” and “threonine-phosphate decarboxylase activity (GO:0048472)” were identified (Table. S3). Totally, ≈ 3 times more categories were identified in the bulk-BEV sequencing, including various metabolic, stress-response, and enzymatic categories (Table. S3). The discrepancy between bulk-BEV sequencing and NP-DS results may indicate a significant difference between genes abundant in quantity and those consistently present in a large proportion of BEVs, highlighting the heterogeneity in DNA encapsulation within BEVs.

### DNA profiling of nanoparticles from human oral biofilm by microdroplet sequencing

We next attempted to characterize DNA content in BEVs produced in the dental plaque biofilm of periodontal patients, the natural habitat of oral pathogenic bacteria. Transmission electron micrographs of the isolated samples revealed that there were spherical particles with diameters ≈ of 100 to 200 nm (Fig. S4A). NTA also detected the particles stained by lipophilic fluorescent dye or double-stranded DNA dye (Fig. S4A), but the number of DNA-staining positive particles was greater than those stained by lipid dye (Fig. S4B), implying the presence of nanoparticles with DNA that does not consist of a lipid layer such as viruses. However, the amount of extracted DNA from those treated samples was below the minimum amount required for conventional metagenome sequencing (< 1 ng per sample).

Remarkably, our droplet sequencing was successful for the collected BEVs with such low amount of DNA. As with the NP-DS for *P. gingivalis*, the sample was mixed with agarose and lipid staining and SYBR-Green-I were used for the number quantification of BEVs and BEVs with amplified DNA (Fig. S2). We could then estimate the fraction of nanoparticles with DNA is ≈ 10 % (Fig. S2BC), which is almost identical to the rate of stained particles by DNA-dye in NTA (Fig. S4B). We next sequenced those 384 positive droplets in total (Fig. 3B). Sequence reads from each droplet were computationally assembled and divided into CDS unit and searched against the nr database (see Materials and Methods). This method detected an average of 52 ± 29 CDSs in one droplet (Fig. 3C), most of which were classified as a virus (bacteriophage)-derived CDS (we call this vCDS hereafter). Of 384 droplets we analyzed, ≈ 20 % of droplets (68/384) contained ≥ 5 bacterial CDSs (we call these bCDSs hereafter) (Fig. 3D), and those droplets were regarded as BEV-containing droplets, and the rest of the droplets contained mostly vCDSs (295/384). When we take into account the fraction of viral particles estimated by the NP-DS, the ratio of BEVs with DNA is estimated as ≈ 30% (Table 1, Fig. S2B, presented as *r*), and thus a significant fraction of BEVs contained DNA in the oral biofilm. We observe particles with the morphology of tailed bacteriophages in TEM (Fig. S5A), all the vCDSs were assigned to *Caudovirales* (Fig. S5B), and thus the origin of the most of viral DNA would be those phage particles. However, we also observed several droplets containing both bCDSs and vCDSs, and the origin of the viral DNA in such cases will be discussed later.

**Figure 3.**
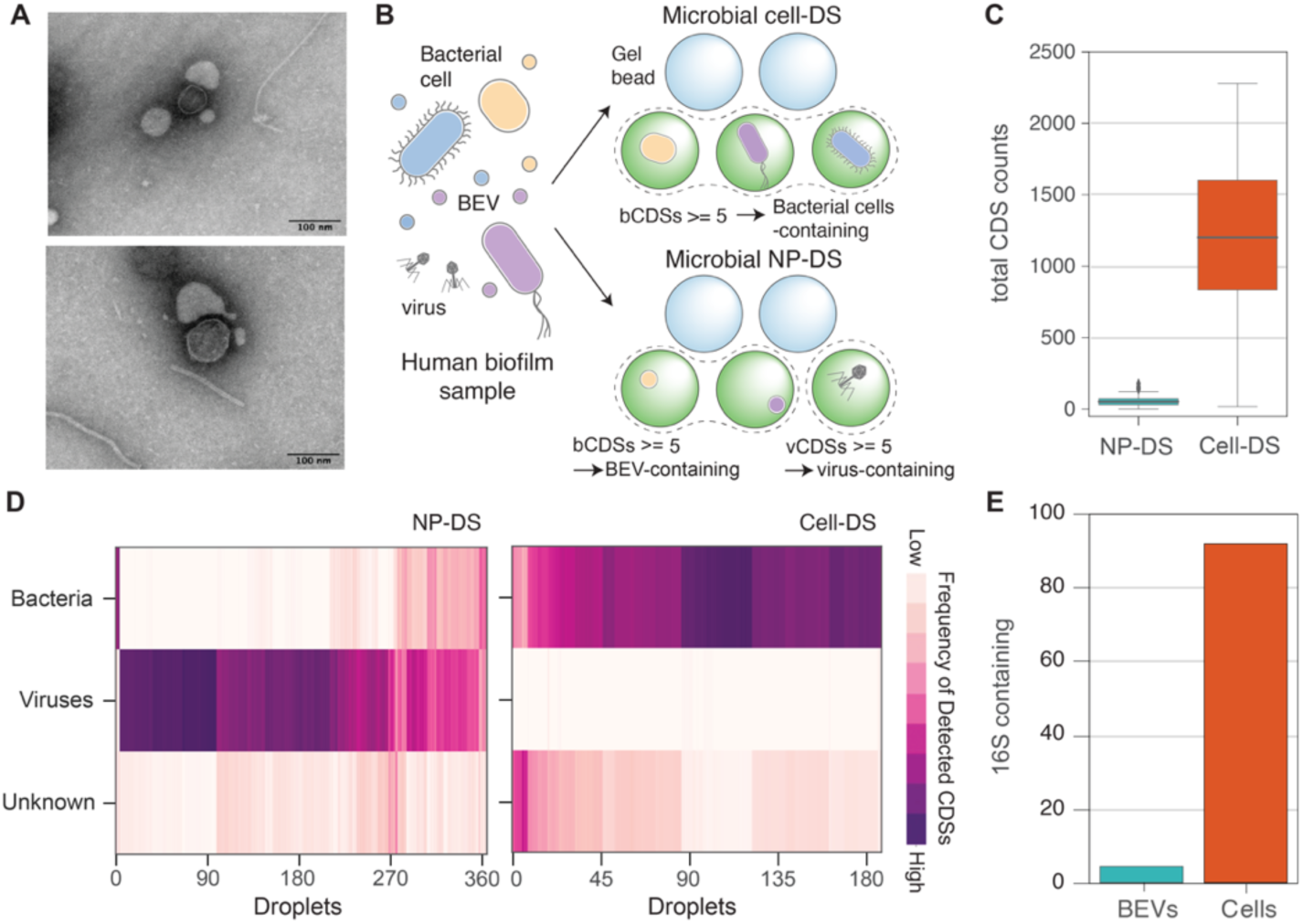
Distinct characteristics of DNA in BEVs compared to bacterial cells. (A) The typical TEM micrographs of BEVs isolated from the dental plaque of periodontal patients. (B) Schematic of microbial cell-DS and NP-DS analysis for the dental plaque biofilm samples from periodontal patients. The separated nanoparticles and bacterial cells in dental biofilms were encapsulated into droplets, and the internal DNA fragments were amplified (see Materials and Methods). In NP-DS, the positive amplification droplets (colored green) were classified by the number of detected bCDSs and vCDSs as BEV-containing, viral DNA-containing, or both. (C) The total length of detected CDSs in NP-DS and cell-DS analysis. The results of all positive droplets from the biofilm samples were analyzed and shown as boxplots. Medians and outliers were shown as bold lines and circles each. (D) Kingdom-level classification of detected CDSs in NP-DS and cell-DS. In each droplet, the length of CDSs assigned to bacteria, viruses, or unknown (i.e., not assigned to any taxon in the database) was normalized by the total length of detected CDSs and shown as a heatmap. The droplets where less than 5 CDSs were detected were eliminated. (E) The percentage of BEV-or bacterial cell-containing droplets in which 16S rRNA sequence was detected. The DNA sample is the mixture of BEVs collected from the biofilm of three patients.

We compared the DNA content in a BEV to that of a bacterial cell by droplet sequencing of bacterial cells (cell-DS) isolated from the same dental plaque samples (Fig. 3B). In cell-DS, averagely 1204 ± 542 CDSs and 790 ± 490 kbp of CDS region was detected (Fig. 3C and Fig. S6A), and full-length 16S rRNA sequences were observed in over 90 % of the droplets (Fig. 3E). In contrast, the CDS regions detected in BEV-containing droplets were mostly around 9 kbp in each droplet (8.6 ± 6.5 kbp, Fig. S6B), corresponding to ≈ 1 % of those detected in cell-DS. Also, full-length 16S rRNA sequences were detected only in 4% (3/68) of BEV-containing droplets (Fig. 3E). The length and prevalence of DNA in BEVs from oral plaque biofilms are consistent with those observed in *P. gingivalis*.

### Enrichment of specific genomic regions in BEVs from the oral cavity

We next taxonomically annotated the detected bCDSs in each droplet by homology search against the GTDB database (Fig. S7). We found that most of the detected bCDSs in each BEV-containing droplet derived from a single phylum, family, and genus-group (around 70 ∼ 90% of purity, Fig. 4A and Fig. S8), and we assumed that the most frequently detected bacterial taxon (MFT) for each droplet reflects an originating bacterium of a BEV. Mapping of the detected sequence reads to the assembly genome of the MFT (Fig. S7) revealed that most of the BEV-containing droplets contained around 0.5 ∼ 1 % of the genomic regions of the MFT, while the droplets in cell-DS reasonably covered an average of 40 ∼ 60 % of the genomic regions of MFT (Fig. 4B).

**Figure 4.**
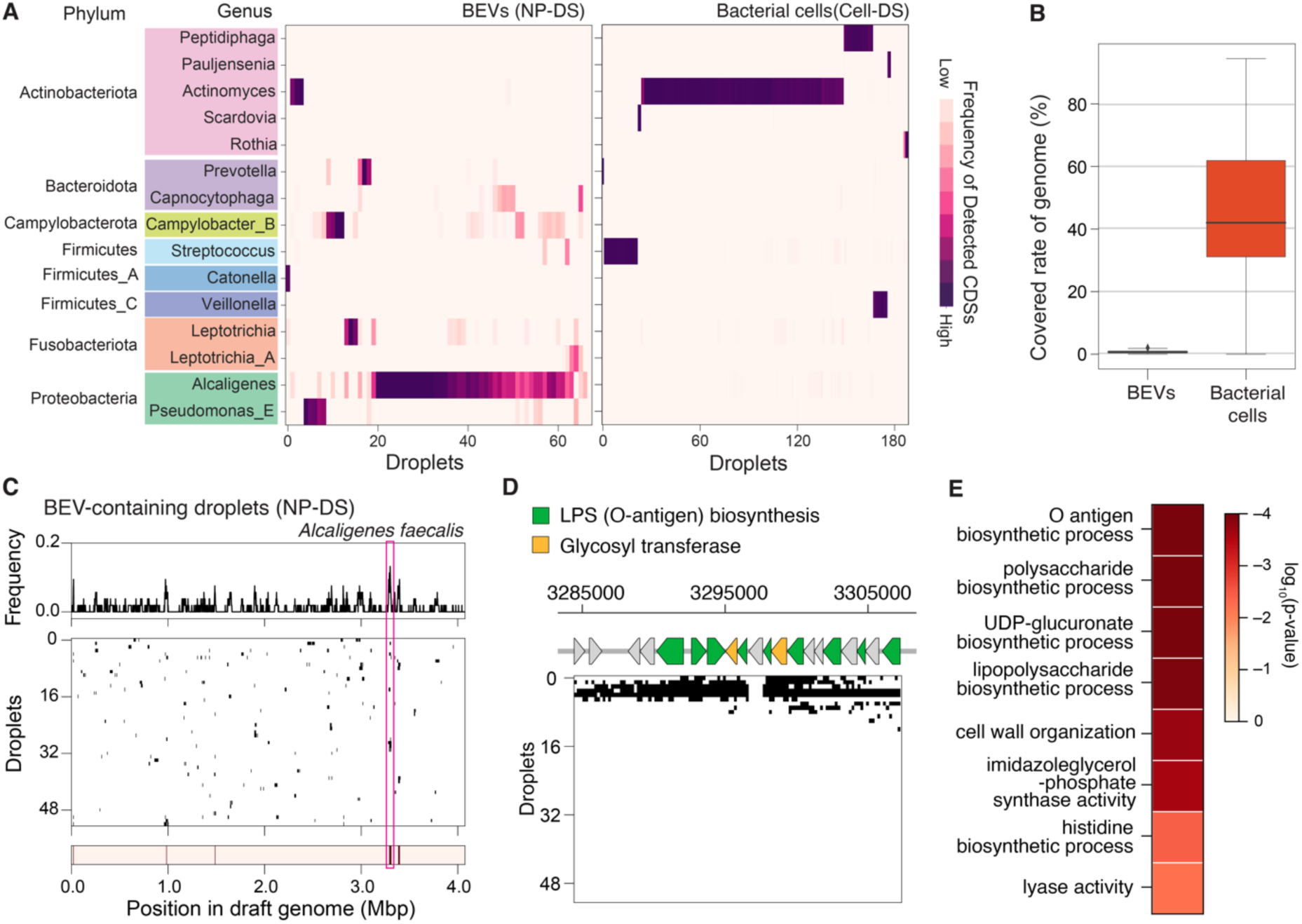
Enriched DNA sequences in BEV-containing droplets. (A) Genus-level taxonomic classification of bCDSs detected in BEV-containing droplets in NP-DS and bacterial cell-containing droplets in cell-DS. The length of CDSs assigned to each bacterial genus group was normalized by the total detected bCDSs and shown as a heatmap. We only showed genera whose maximum frequency of detected CDSs in a single droplet is more than 0.4. (B) The percentage of the genomic region covered by the sequence reads in a BEV-or bacterial cell-containing droplet. For each droplet, sequence reads were aligned to the assembled genome of the MFT. Medians and outliers were shown as bold lines and circles each in box plots. (C) Genomic regions in *Alcaligenes faecalis* (GCF_000745965.1) mapped by sequence reads in BEV-containing droplets. The frequency of droplets including each genomic region (top). The positions on the genome where the sequence reads were mapped in each droplet were filled with black (middle), The genomic regions which were significantly enriched in BEV-containing droplets were filled with red (bottom). We analyzed sequence data of 53 BEV-containing droplets whose MFT was determined as *Alcaligenes faecalis* (GCF_000745965.1). (D) Detection profiles of the LPS biosynthesis gene cluster (encircled by magenta in panel C) in 53 droplets. Green gene arrows correspond to genes associated with LPS (O-antigen) biosynthesis. Yellow gene arrows correspond to glycosyl transferase genes, which are also possibly related to O-antigen biosynthesis process. (E) Screened functional categories (GO terms) as that were significantly overrepresented in the enriched genomic region of BEVs from *Alcaligenes faecalis*. Same as Fig. 1F, we statistically extracted the GO terms (hypergeometric test, p-value < 0.05, see Supporting Materials and Methods). The statistical significance level is shown as a heatmap.

The enrichment of specific genomic regions was also observed in BEVs from the dental plaque biofilm as with pure culture system. We focused on droplets whose MFT is *Alcaligenes faecalis* (GCF_002443155.1), which is most frequently observed taxon as MFT in the BEV-containing droplets (Fig. 4C). The mapped regions by sequence reads differed among droplets and distributed to the wide range of the host chromosomal region (Fig. 4C). However, there were commonly detected regions among the droplet (Fig. 4C, binominal test, p-value < 0.01). Among the 46 CDSs located on the enriched regions, many of the genes related to lipopolysaccharide (LPS) biosynthesis (Table S4), and most of those are clustered in one genomic locus (Fig. 4D). A statistical screening of enriched functional categories (GO terms) in those genes revealed that most of the screened GO terms are associated with biosynthesis of bacterial antigen (LPS), such as “O antigen biosynthetic process (GO:0009243)” and “LPS biosynthetic process (GO:0009103)” (Fig. 4E and Table S5). “O-antigen biosynthesis genes” were encapsulated in ≈ 14 % of BEVs of *Alcaligenes faecalis* (≈ 10 % of all analyzed DNA-containing BEVs in the oral biofilm), suggesting the prevalence of these gene in the biofilm BEVs.

### Origins of BEV-derived DNA in dental plaque

Taxonomic profiles of bCDSs detected in NP-DS showed quite different compositions from those detected in cell-DS. In phylum-level, abundantly detected groups in BEV-containing droplets such as *Proteobacteria* were barely detected in cell-DS (Fig. 4A), and thus most of the DNA sequences detected in BEVs derived from bacterial species with very low abundance in dental plaque biofilm. Indeed, *Actinomyces, Peptidiphaga,* and *Streptococcus* were the most frequently detected taxa and MFTs in ≈ 90 % of droplets in cell-DS, while those CDSs were barely detected in NP-DS (only in 6 droplets in total) (Fig. 4A). On the other hand, CDSs from *Alcaligenes* (*Proteobacteria*), the most abundant genus detected in BEV-derived DNA, were quite low in frequency in cell-DS (Fig. 4A). In addition to *Alcaligenes*, we also found the bCDSs from several minor bacterial genus in cell-DS, such as *Prevoteolla, Capnocytophaga, Campylobacter_B, Leptotrichia, and Pseudomonas_E*, in NP-DS. Those results suggest that the detected BEV-derived DNA by our method could reveal the distribution of functional genes among particles, but also highlight the taxonomically minor but active producer strains of BEVs in microbiome.

## Discussion

Understanding the diversity of DNA cargo within BEVs and identifying genes consistently present is crucial for comprehending the role of HGT at the microbiota level. Traditional metagenomic approaches, however, have struggled to accurately characterize such properties. Our study addresses this limitation by employing droplet sequencing analysis, a method that reveals the overlooked complexity of DNA content in BEVs. We found that BEVs typically harbor substantial DNA fragments (ranging from 10 to 50 kb) from the host genome, with certain regions being prevalent across the BEV population, and those enriched genomic regions contain functionally related gene clusters (Fig. 2, and Fig. 4). Our analysis demonstrates a substantial refinement in identifying functionally relevant genes within BEVs using NP-DS, compared to bulk-BEV sequencing (Fig. 1). The latter tends to capture a broader array of miscellaneous gene sets (as detailed in Data S1) and is susceptible to amplification biases during library preparation. It is possible that DNA amplification within individual droplets, combined with count-based statistical analysis, mitigates these biases, leading to a more precise identification of functionally enriched genes within BEVs. Furthermore, the sensitivity of our technique to detect minute quantities of DNA expands the potential of NP-DS for broader application across a variety of microbiome samples.

Enrichment profiles and prevalence of DNA content suggest their potential ecological impacts on the microbial community via HGT. The enrichment of a specific group of metabolic or virulence-associated genes was found in both bacterial cases (Fig. 2A and Fig. 4E), which is reminiscent of similarities with other mobile genetic elements (e.g., plasmid and integrative conjugative element) (33, 34). For both cases, clustered gene sets are found to be 3 - 12 % of BEVs, which is a hundred times greater than what was detected in the natural environment using traditional DNA dye staining techniques (22), revealing DNA-packaging BEVs are far more prevalent than previously estimated.

“O-anitgen biosynthesis genes” were quite prevalent in the BEVs derived from the oral plaque biofilm. This gene sets are generally clustered on the genome of gram-negative bacteria (35, 36), and high sequence diversity and unique GC content of that genomic region suggests that this gene cluster is a frequent site of HGT (37, 38). Given that divergence in O-antigen have an potential impact on the host-bacterial interactions (39), one possibility is that the enriched genes in BEVs could modulate pathogenicity of the oral biofilm via HGT.

In BEVs from *Porphyromonas gingivalis*, we observed the enrichment of genes associated with DNA integration, such as those involved in DNA transposition and CRISPR-Cas systems. The interspecies exchange and transposition of insertion sequences (IS) are believed to contribute to genome rearrangements in this pathogen (40). Intriguingly, the CRISPR spacer sequences are highly homologous to regions of IS on the *P. gingivalis* genome, and those spacer sequences are possibly responsible for preventing IS transposition and recombination (40). The enrichment of both IS and CRISPR-Cas elements suggests that BEVs in this bacterium play an important role in the interspecies recombination of IS and its control by the CRISPR-Cas system by mediating the HGT. While the frequency and extent to which these genes are transferred among bacteria warrant further investigation, our study lays the groundwork by identifying transferable gene candidates via BEVs and their potential effects on microbial community dynamics.

Our findings not only highlight the enrichment of the specific genomic locus in BEVs but also revealed considerable variation in the profiles of BEV-containing droplets (Fig. 1B and Fig. 4D), suggesting that the packaging process influenced by both probabilistic and deterministic factors. For *P. gingivalis,* the BEV DNA is notably rich in transposable elements, suggesting that these DNA regions tend to be excised and loaded into the BEVs via certain mechanisms. While the excision of the IS element itself is generally considered as a stochastic event, it can be influenced by environmental stressors or specific genetic factors (41, 42), which may alter the frequency of their incorporation into BEVs (43).

Conversely, in other genomic loci where we did not detect signatures of excision, the inclusion of specific DNA fragments into BEVs might not be driven by site-specific excision. In such a case, fragmentation of chromosomal DNA upon explosive cell lysis would be one plausible mechanism for generating small pieces of DNA. The mechanism underlying selective DNA packaging into BEVs may vary; for instance, *Janas et al.* proposed the difference in affinity of an RNA molecule to the membrane as a mechanism for a high abundance of the specific RNA (44). A similar principle could apply to DNA, where molecular affinities and interactions influence the selection process for vesicle encapsulation.

While prior research has attempted to identify BEV producers using 16S rRNA sequencing, our findings underscore the rarity of 16S rRNA sequences within BEVs (refer to Fig. 1E and Fig. 3E), highlighting the limitations of 16S rRNA-based taxonomic identification, such as amplicon sequencing. Instead, our droplet-based sequencing method provides a more comprehensive view of the taxonomic composition of BEV-derived DNA, revealing substantial taxonomic difference between the profiles obtained from cell-DS and NP-DS (Fig. 4A). This discrepancy suggests that the minor members more actively produce BEVs or package DNA into BEVs than the major bacterial group in the human dental plaque. NP-DS identified DNA from several other pathogens (e.g., *Capnocytophaga* and *Leptotrichia*), which were quite minor bacterial groups in our cell-DS data (Fig. 4A), but some of those were prominent pathogens in periodontitis (45). Assuming controlled DNA encapsulation into BEV is the reflection of microbial activity, these enriched DNA regions may serve as biomarkers for assessing the activity of such pathogens. Therefore, the unique characteristics of BEV-derived DNA imply their potential applicability for biopsy in diagnosis.

Our analysis also detected much of phage-derived DNA in NP-DS. Although the origin of those viral DNA fragments would be mostly phage themselves, around 20 % of droplets contained both bacterial and viral CDSs (i.e., more than 5 bCDSs and vCDSs). Given that the samples were treated with DNase I to remove externally existing DNA, it’s plausible that these viral fragments were incorporated into BEVs during phage infection or induction, (21, 46) or by phages targeting BEVs as ‘decoy’ cells (47). Upon identifying key virulent BEV producers and associated phages, such DNA insertion into BEVs has the potential to be harnessed for regulating HGT in phage therapy applications. While further experimental investigation is necessary to understand the coexistence of bacterial and phage DNA, the droplet sequencing would be a potential tool to explore potential phage-BEV interactions in nature.

## Materials and Methods

### Isolation and purification of BEVs from *P. gingivalis*

*P. gingivalis* W83 was grown in Gifu anaerobic medium at 37°C. To maintain anaerobic conditions, 20 min of N_2_/CO_2_ (80:20 v/v) gas sparging was performed prior to culture inoculation. The culture was grown at 37 °C until the OD_600_ reached 1.0. Then, the culture was centrifuged for 10 min at 7800 rpm and 4 °C. The resulting supernatant was passed through 0.22 µm filters to remove any cell debris. The filtered supernatant was ultracentrifuged for 2 h at 126,000 × g at 4 °C and the resulting pellets were re-suspended in phosphate buffered saline (PBS).

### DNase I treatment

DNase I (13 units (U) / µL) (NIPPON GENE) was added to a final concentration of 2U to treat the external DNA of nanoparticles. This sample was stored at 37 °C for 30 minutes, followed by DNase I deactivation by heating the sample at 80 °C for 10 minutes.

### DNA sequencing of bulk BEV samples

We purified DNA from DNase I-treated BEV samples using the Isoplant-II DNA extraction kit (NIPPON GENE) according to the manufacturer’s instructions. Sequencing libraries were prepared using the QIAseq FX DNA Library Kit (QIAGEN) according to the manufacturer’s protocols and sequenced using the Illumina Nextseq 2000 with 2 × 150 bp configuration. The obtained fastq files were processed using fastp 0.19.5 (48), and the sequence reads were aligned to the assembly genomes of the host bacterium (*P. gingivalis*: GCF_000007585.1) by bowtie2 (49) with option (--sensitive-local).

### Nanoparticle tracking analysis (NTA)

NTA was performed by ZetaView (Particle Metrix) for the quantification of nanoparticles. Samples were measured by scanning 11 cell positions and capturing 30 frames per second. For measurements using fluorescent signals, 488/500 nm and 660/680 nm laser-filter units were used with the camera setting: Sensitivity: 70 ∼ 80; Shutter: 200; Minimum trace length: 15. For measurements in scatter mode, we used the following settings: Sensitivity: 65; Shutter: 200; Minimum trace length: 15. Captured video images were further analyzed using the ZetaView Software.

### Droplet genome sequencing

Droplet genome sequencing was performed by bitBiome (Japan) as previously described (50), and details were described in Supporting Materials and Methods. Briefly, the cell or nanoparticle suspensions were mixed with 1.5 % agarose solutions so that the expected percentage of positive droplets was ≈ 40 % (23). In the case of cell-DS, the encapsulated cells were further lysed by the lysis solutions in gel beads. The droplets in cell-DS and NP-DS were processed for multiple displacement amplification (MDA) by REPLI-g Single cell Kit (QIAGEN), and the beads were stained with 1× SYBR Green (Thermo Fisher Scientific). Green fluorescence-positive beads were sorted by the FACSMelody cell sorter (BD Bioscience). The collected positive droplets were proceeded to the second round of WGA with the REPLI-g Single Cell Kit. The whole-genome amplified samples with enough DNA were further subjected to the whole-genome sequencing analysis using the Nextera XT DNA Library Prep Kit (Illumina) according to the manufacturer’s protocols and sequenced using the Illumina HiSeq.

### Acquisition of transmission electron micrographs (TEM)

Purified BEV samples were placed onto formvar coated copper grids and negatively stained with 2% phosphotungstate acid solution (pH = 7.0) for 60 sec. The grids were analyzed and visualized using JEM-1400 microscope (JEOL Ltd.) operated at an acceleration voltage of 100 kV and imaged by EM-14830RUBY2 CCD camera (JEOL Ltd.).

### Subjects and dental plaque sample collection for droplet sequencing

We recruited patients with periodontitis, who were classified as generalized stage III-grade C (51). These patients had not taken antimicrobials within 3 months, were non-smokers, and did not have diabetes or any serious systemic disease. The present study was approved by the ethical committee at Kyushu Dental University (ethical approval number: 26-28) and conducted in accordance with the approved guidelines. All participants signed informed consent.

The collection of dental plaque biofilm samples (n = 3) was performed based as follows. Rolled cotton was placed next to the teeth to prevent saliva contamination. Supra- and subgingival plaque was collected by a periodontist using a curette without bleeding. The collected dental plaque biofilm sample was immediately resuspended into 1 mL PBS. Then the biofilm was treated with 100 µg/mL Proteinase K (QIAGEN) for 1 hour at 37 °C with vortexing at 20-minutes intervals to degrade extracellular matrices, and the treated sample was centrifuged for 10 minutes at 6000 g at room temperature. The resulting pellet was resuspended into OMNIgene^®^-ORAL OM-501 (DNA Genotek) and used for cell-DS. The supernatant was passed through 0.22 µm filters to remove any extracellular substrates. The filtered supernatant was ultracentrifuged for 2 h at 200,000 × g at 4 °C, and the resulting pellets were re-suspended in 10 mM HEPES with 0.85 % NaCl.

### Data analysis in NP-DS and cell-DS

The paired-end read sequences from each droplet were first processed by fastp 0.19.5 and then assembled by SPAdes 3.12.0. The viral contigs were extracted by VIBRANT 1.2.1 (52) to extract viral contigs. Then coding sequences (CDSs), rRNAs, and tRNAs were extracted from the contigs by Prokka 1.14.6 (53). The extracted CDSs were grouped together and clustered by CD-HIT. The clustered CDSs were first annotated by homology search against the National Center for Biotechnology Information (NCBI) non-redundant (nr) database (downloaded on July 1^st^, 2021) by DIAMOND 2.0.8 (54). For each CDS, the Genbank accession number of the best-hit protein sequence was further used for taxonomic annotation by BASTA (55). The CDSs classified as Bacteria in the kingdom-level were further annotated by homology search against protein sequences in the Genome Taxonomy Database (GTDB) (56) (version r202) by DIAMOND for more strict taxonomic annotation of bacterial CDSs with a threshold in a percent identity ≥ 80 %. Then we eliminated bacterial taxa that were classified as potential contaminants in low-biomass human samples in previous studies (57, 58) (see Data S2). For droplets determined to BEV-containing, we first grouped CDSs by GTDB accession number and calculated the total CDS length assigned to each GTDB taxonomy. Then the most abundantly detected GTDB taxonomy is designated to the “most frequently detected taxon” (MFT) for each droplet. For the details of computational analysis, see the Supporting Materials and Methods.

## Supporting Information

Supporting Materials and Methods Figs. S1-S8

Tables S1-S5

Datasets S1-S2

## Acknowledgments

This work was financially supported by JST, PRESTO (JPMJPR19H1 to A.O.), a Grant-in-Aid for Research from the Japan Society for the Promotion of Science KAKENHI (grant numbers 22H02265 to A.O.), and the Japan Agency for Medical Research and Development (19gm6010002h0004 to A.O. and 21ae0121044h0001 to A.O. and So.T.).

## Author contributions

So.T. and A.O. designed the research; So.T. developed analysis pipelines and analyzed the data; N.D., Sa.T., Y.K., T.M., M.U., and A. W. performed the experiments; So.T. and A.O. wrote the final version of the paper; All authors contributed to data interpretation and the writing of the manuscript.

## Data availability

All sequence raw data used in this study were deposited in the DNA Data Bank of Japan (DDBJ) with the accession ID PRJDB17260 and PRJDB17266.

## Supporting Materials and Methods

### Droplet genome sequencing

Droplet genome sequencing was performed by bitBiome (Japan) as previously described (1). Briefly, in the cell-DS, the cell suspensions from biofilm samples were mixed with 1.5 % agarose solutions in Dulbecco’s PBS (DPBS; Thermo Fisher Scientific) so that the expected percentage of positive droplets was ≈ 40 % (i.e. cells/droplets ratio become ≈ 0.5) based on the previous study (2). In the case of nanoparticle genome sequencing, the suspensions of nanoparticles were mixed with the agarose solutions so that the expected percentage of positive droplets was less than 30 %. Bacterial cells or nanoparticles were encapsulated in droplets using fabricated microfluidic droplet generators (3) and further used for in-bead bacterial genome amplification sequencing. Collected droplets were incubated on ice for 15 min for solidification. The solidified gel droplets were broken with 1H,1H,2H,2H-perfluoro-1-octanol (Sigma-Aldrich). Then, the gel beads were washed with acetone (Sigma-Aldrich), and the solution was mixed vigorously and centrifuged. The gel beads were washed three times each with DPBS, followed by isopropanol. In the case of cell-DS, the encapsulated cells were further lysed by the lysis solutions in gel beads. First, 50 U/ μL Ready-Lyse Lysozyme Solution (Epicentre), 2 U/ mL Zymolyase (Zymo Research), 22 U/mL lysostaphin (MERCK), and 250 U/mL mutanolysin (MERCK) were added and incubated in DPBS at 37 °C overnight. Then gel beads were washed three times and 0.5 mg/mL achromopeptidase (FUJIFILM Wako Chemicals) was added into DPBS at 37 °C for 8 hours. The gel beads were washed with DPBS three times again and 1 mg/mL Proteinase K (Promega) with 0.5% SDS in PBS was added and incubated overnight at 40 °C. Following lysis, the gel beads were washed with DPBS five times. Then, the droplets in cell-DS and NP-DS were processed for multiple displacement amplification (MDA) by REPLI-g Single cell Kit (QIAGEN). The droplets were suspended by Buffer D2 from a REPLI-g Single Cell Kit and incubated for 2 hours or 8 hours for cell-DS or NP-DS, respectively. Following whole-genome amplification (WGA), gel beads were washed three times with 500 μL DPBS. Thereafter, beads were stained with 1× SYBR Green (Thermo Fisher Scientific) in DPBS. Following the confirmation of DNA amplification by the presence of green fluorescence in the gel, fluorescence-positive beads were sorted into 0.8 μL DPBS in 96-well plates using the FACSMelody cell sorter (BD Bioscience) equipped with a 488-nm excitation laser. Following droplet sorting, 96-well plates were proceeded to the second round of WGA. Second-round MDA was performed with the REPLI-g Single Cell Kit. The 96-well plates were incubated at 65 °C for 10 min after the addition of Buffer D2 (0.6 μL) to each well. Thereafter, 8.6 μL of MDA mixture was added to each well, and the plates were incubated at 30 °C for 2 hours for cell-DS and 8 hours for NP-DS, followed by the incubation at 65 °C for 3 minutes. Following second-round amplification, aliquots of the 96 well-plates were transferred to replica plates for DNA yield quantification using the Qubit dsDNA High Sensitivity Assay Kit (Thermo Fisher Scientific). For cell-DS, 16S rRNA gene Sanger sequencing (FASMAC) with the V3-V4 primers (Forward, 5′-TCGTCGGCAGCGTCAGATGTGTATAA GAGACAGCCTACGGGNGGCWGCAG-3′; reverse, 5′-GTCTCGTGGGCTCGGAGATGTGTATAAGAGAC AGGACTACHVGGGTATCTAATCC-3′) was also performed using the replica plates. The whole-genome amplified samples with enough DNA were further subjected to the whole-genome sequencing analysis. In the case of cell-DS, the whole-genome amplified samples were further screened using the Geneious software (Biomatters, Ltd.) based on the quality of 16S rRNA with the criteria that more than 400 bp in length and HQ 40%. For whole-genome sequencing analysis, sequencing libraries were prepared from second-round MDA products using the Nextera XT DNA Library Prep Kit (Illumina) according to the manufacturer’s protocols and sequenced using the Illumina HiSeq 2 × 150 bp configuration for the biofilm samples or 2 × 75 bp configuration for *P. gingivalis* BEVs.

### Estimation of the percentage of particles with DNA in cell-DS and NP-DS

The theoretical ratio of droplets to particles in the encapsulation step is denoted as *Rt* and described as follows:

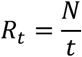

Here *N* is the total number of nanoparticles (cells) and *t* is the number of droplets. We used the fixed *Rt* for cell-DS or NP-DS as described above. Here, the nanoparticle samples subjected to the gel-lysis are considered as a mixture of viruses, BEVs, and other nanoparticles (extracellular proteins), and thus *N* can be described as follows:

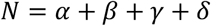

Here, *α*, *β*, *γ*, and *δ* are the total numbers of viruses, BEVs with DNA, BEVs without DNA, and other nanoparticles. The observed ratio of droplets to particles with DNA in the encapsulation step is denoted as *Ro* and described as follows:

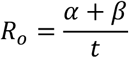

For estimating *Ro*, the percentage of DNA-containing droplets (we call this parameter as *Po* hereafter) after processing cells or nanoparticles were quantified by counting the number of droplets with green fluorescence in microscopic images. From *Po*, we further estimated the ratio of droplets to particles with DNA (*Ro*) using the empirical curve in a previous study (2). We can also estimate the ratio of BEVs (*Rb*) by positive particles by lipid-staining dye using NTA.

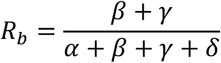

The ratio of droplets containing viruses to those containing BEVs with DNA was estimated based on the taxonomic profiles (Fig. 3).

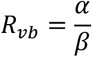

Then the percentage of BEVs with DNA (*r*) was calculated as follows.

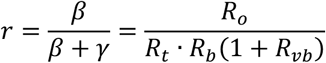

### Pre-processing of sequence reads and extraction of CDSs in droplet sequencing

The paired-end read sequences from each droplet were first processed by fastp 0.19.5 (4) for quality control and adapter trimming. The processed sequence reads were assembled by SPAdes 3.12.0 (5) with options (-k auto –disable-rr--careful). The generated contig was used for further processing by VIBRANT 1.2.1 (6) to extract viral contigs. Then CDSs, rRNAs, and tRNAs were extracted from the contigs by Prokka 1.14.6 (7).

### Taxonomic annotation of CDSs in droplet sequencing

The extracted CDSs from droplets derived from the same sample were grouped together and clustered by CD-HIT (8) with options (-c 0.98 -s 0.5 -aS 0.9). The clustered CDSs were first annotated by homology search against the National Center for Biotechnology Information (NCBI) non-redundant (nr) database (downloaded on July 1^st^, 2021) by DIAMOND 2.0.8 (9) with options (--evalue 1e-10 –outfmt 6 –sensitive). For each CDS, the Genbank accession number of the best-hit protein sequence was further used for taxonomic annotation by BASTA (10) with options (sequence prot −l 100 -I 80 -b). At this step, hits whose percent identity is less than 80 % and whose matched length is less than 100 bp were discarded. Genbank accession numbers that were not mapped to the default BASTA database were further mapped to the NCBI database using custom python code implemented in the original BASTA pipeline. Briefly, taxonomic information of each protein sequence in DIAMOND hits was obtained by Bio.Entrez module in Biopython package (11) and formatted by TaxonKit (12).

The CDSs classified as Bacteria in the kingdom-level were further annotated by homology search against protein sequences in the Genome Taxonomy Database (GTDB) (13) (version r202) by DIAMOND for more strict taxonomic annotation of bacterial CDSs. If a top-hit of CDS to the GTDB protein database with a percent identity ≥ 80 %, the assembly accession number registered in GTDB was assigned. We eliminated bacterial taxa that were classified as potential contaminants in low-biomass human samples in previous studies (14, 15). The list of eliminated GTDB taxa is shown in Data. S2.

### Screening of most frequently detected taxon in BEV-containing droplets

For droplets determined to BEV-containing, we first grouped CDSs by their taxonomic information (GTDB accession number) and calculated the total CDS length assigned to each GTDB taxonomy. Then the most abundantly detected GTDB taxonomy in the total CDS length is designated to the “most frequently detected taxon” (MFT) for each droplet.

### Screening of enriched genomic regions in BEVs from dental plaque biofilm

We first aligned sequence reads obtained from each droplet to the assembly genomes of host bacteria by bowtie2 (16) with option (--sensitive-local). We analyzed BEV-containing droplets from the *P. gingivalis* and mapped the sequence reads to the host bacterial genome (*P. gingivalis*: GCF_000007585.1). It is well known that MDA usually harbors chimeric sequences during the amplification process (17) and it is possible that presence of those chimeric reads affect the mapping profiles. We checked this possibility by BWA (18) and found that only 1.41 ± 0.34 % of reads were classified as chimeric in 96 droplets. We also confirmed that cleaning of those chimeric reads by recently developed algorithm (19) did not harbor qualitative difference in the mapping profiles compared to those generated by the original read sequences (Fig. S3). In the case of BEV-containing droplets in the dental biofilm samples, we focused on *Alcaligenes faecalis* (GCF_002443155.1), which were frequently detected as MFT in droplets. Then, the assembly genome sequences for each bacterium were separated into 1000 bp sections, and significantly enriched genomic regions in the droplets were screened by a binominal test with the following formula:

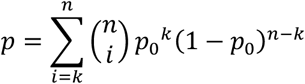

Where *n* is the total number of droplets examined, *k* is the number of positive droplets (i.e. droplets in which the targeted 1000 bp section was detected), and *p0* is the average frequency of positive droplets for the overall genomic region. If *p* (p-value) is less than 0.01, that 1000 bp section is regarded as an enriched genomic region in BEV-containing droplets.

### Gene enrichment analysis in BEV-containing droplets

The significantly enriched genomic regions were divided into CDS-unit by Prokka and functionally annotated by homology search using DIAMOND against UniprotKB/Swiss-Prot database (20) (Downloaded on April 21, 2021). To see the enrichment of specific functional categories in BEV-derived CDSs, the gene ontology (GO) annotations (21) are attributed to a top hit in UniprotKB/Swiss-Prot database assigned to each CDS. Then enriched GO annotations in BEV-derived CDSs were screened by a hypergeometric test with the following formula:

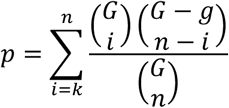

Where *n* is the total number of enriched CDSs, *G* is the total number of CDSs in the genome, and *g* is the total number of CDSs in a GO annotation category of interest. The probability that more than *k* CDSs were found in each GO annotation category is denoted as *p*. If *p* is less than 0.05, that GO annotation is regarded as an enriched functional category in BEV-containing droplets.

---

**Figure S1.**
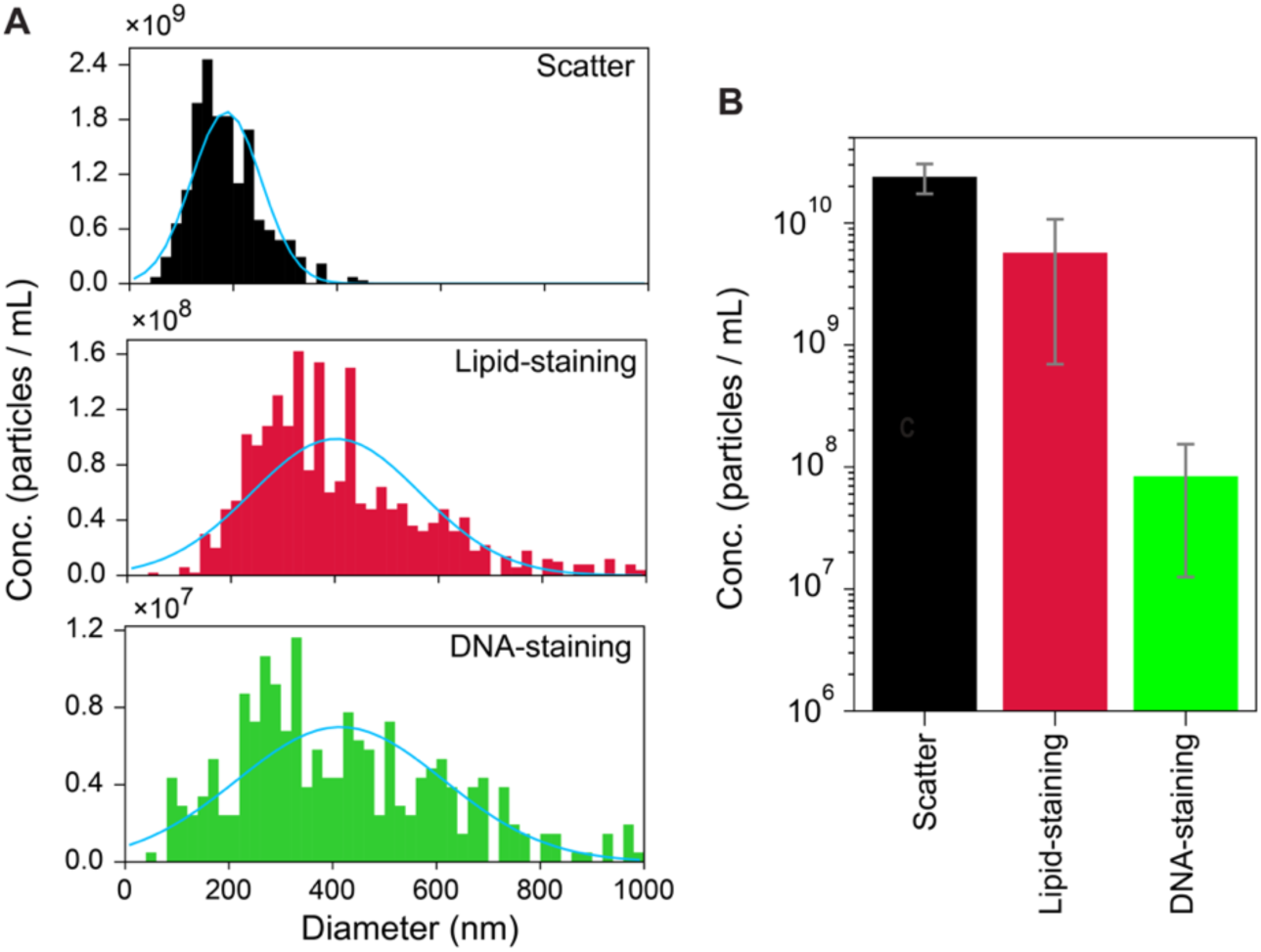
NTA (nanoparticle tracking analysis) of BEVs isolated from pure cultures of *P. gingivalis*. (A) The typical size distribution of nanoparticles isolated from pure cultures of *P.gingivalis* by NTA (nanoparticle tracking analysis). We stained the isolated nanoparticles with the lipid-staining dye, 33.3 µM DiD (1,1’-Dioctadecyl-3,3,3’,3’-Tetramethylindodicarbocyanine), and DNA-staining dye, 2.67 µg/mL 1000×SYBR-Green I. We detected the scattered light signals (Scatter, black), lipid dye fluorescence (red, detected in 660/680 nm laser-filter unit), or DNA dye fluorescence (green, detected in 488/500 nm laser-filter unit). A blue line is the fitted curve of the normal distribution to the experimental data. (B) The total concentration of nanoparticles detected by each signal with 50 ∼ 500 nm in size. Error bars indicate the standard deviations in triplicate experiments.

**Figure S2.**
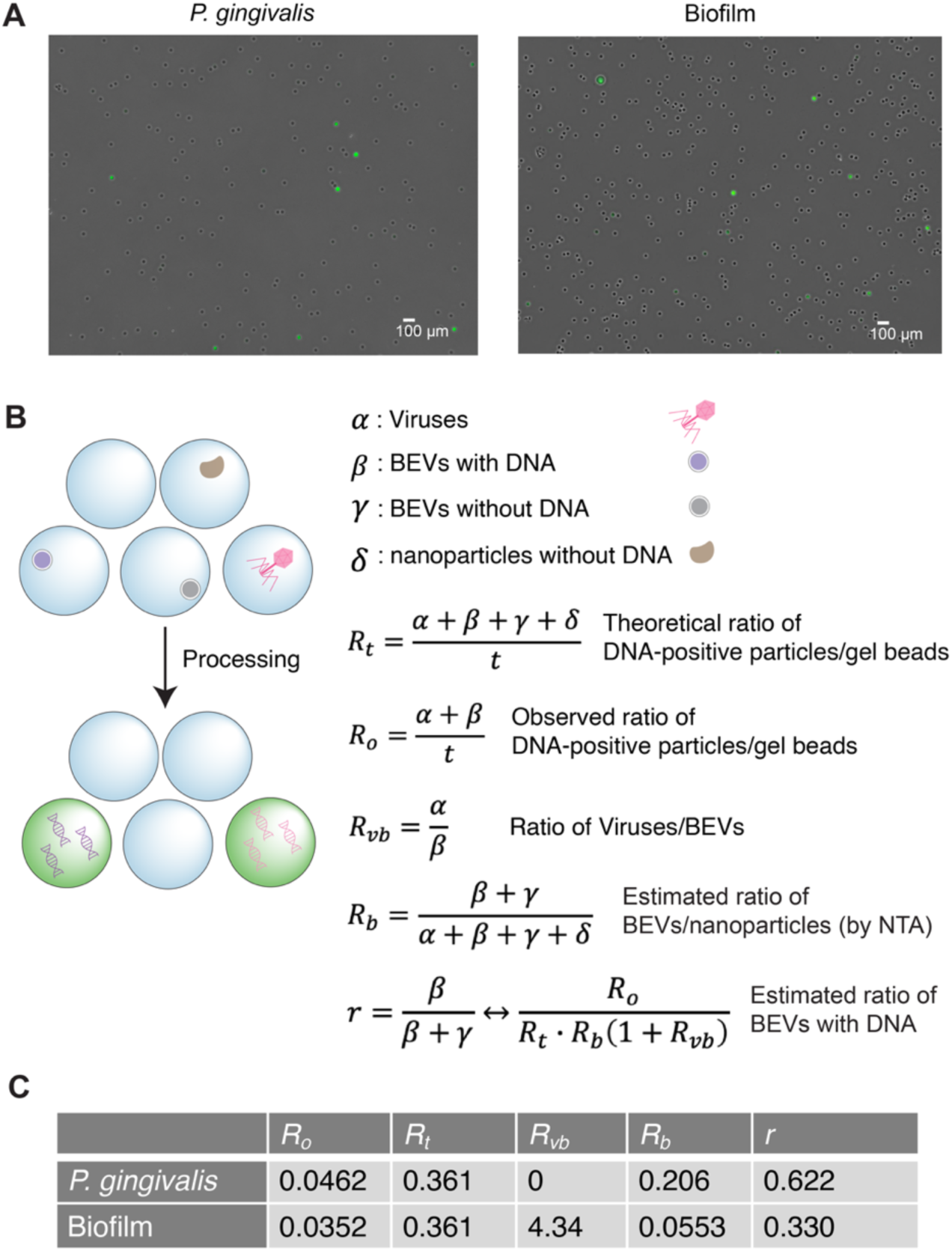
The percentage of droplets containing amplified DNA fragments after processing cells or nanoparticles. (A) Representative microscopic fluorescence images of gel beads after in-gel lysis and amplification. The beads were stained by SYBR-Green-I to check the number of positive droplets (i.e., internal DNA was successfully amplified). (B) Schematic of estimating the percentage of nanoparticles with DNA. The nanoparticles expected to contain DNA, BEVs with DNA or phages, were colored by purple and red, respectively. Gel beads expected to contain amplified DNA fragments were colored green. The theoretical ratio of droplets to total particles (*Rt*) was controlled to a fixed value in our analysis (see Supporting Materials and Methods). The ratio of droplets to particles with DNA (*Ro*) was estimated by the percentage of positive droplets in a microscopic image and an empirical curve in a previous study. Based on those two parameters and other two parameters estimated by NTA and taxonomic annotation (see Supporting Materials and Methods), we calculated the percentage of BEVs with DNA, *r*. (C) The observed percentage of positive droplets and estimated percentage of BEVs with DNA.

**Figure S3.**
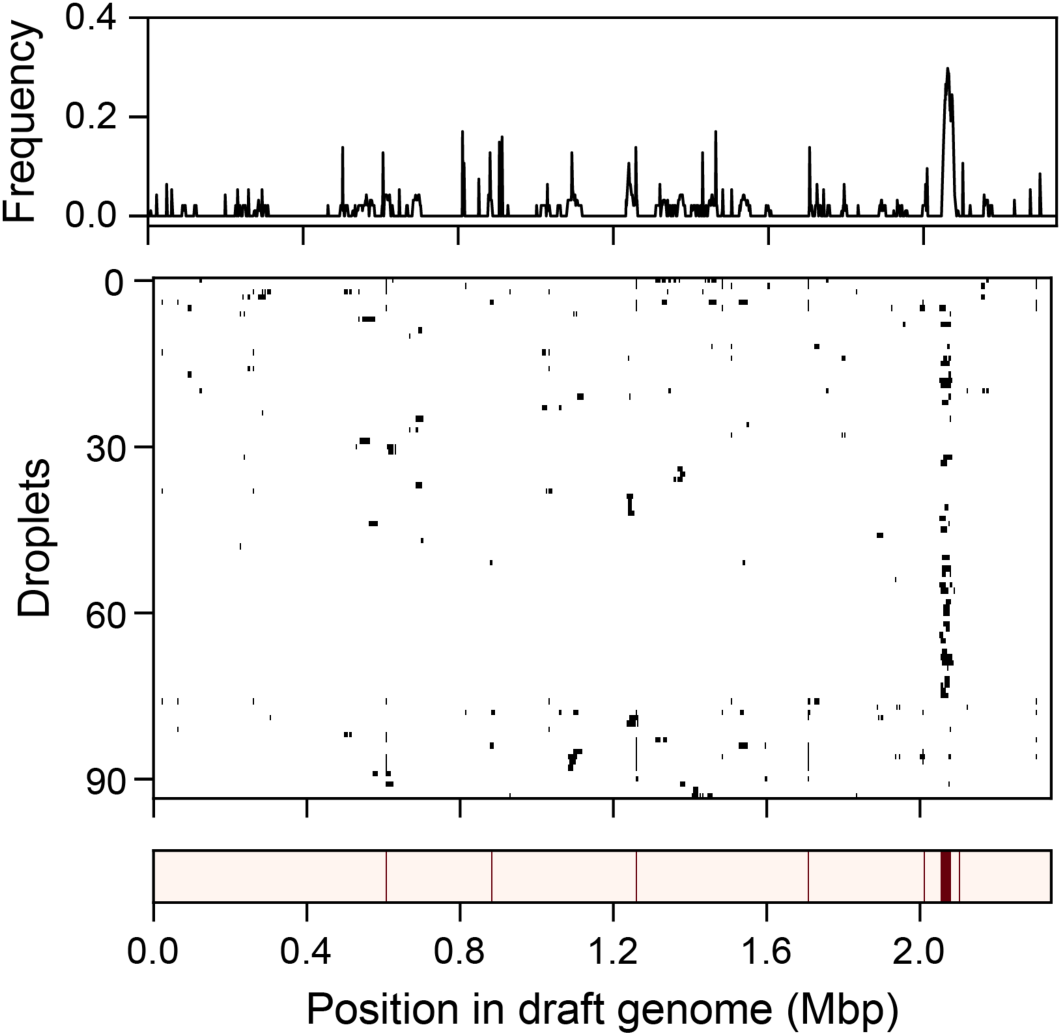
Mapping profiles of sequence reads from *P. gingivalis* BEVs after cleaning of chimeric reads. The sequence reads were mapped after the cleaning process of chimeric reads by bwa. Same as Fig. 1B, the genomic positions where the sequence reads were mapped in each droplet were filled with black.

**Figure S4.**
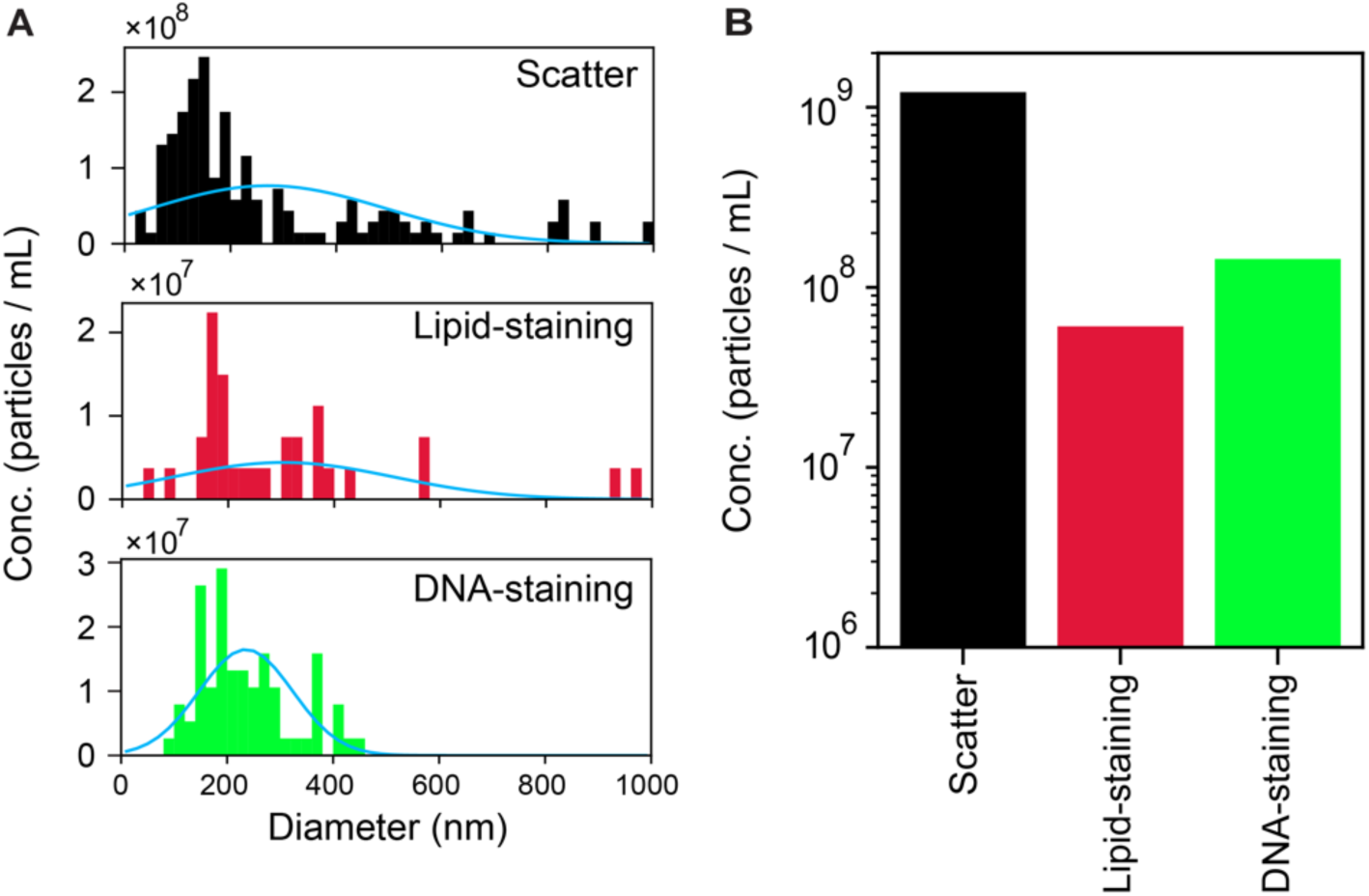
Characterization of isolated nanoparticles from the dental plaque biofilm. (A) The size distribution of nanoparticles isolated from the dental plaque sample. The samples were stained by 5 µg/mL lipid staining FM4-64 for and 1 µg/mL of 1000×SYBR Green I. We detected the scattered light signal (black), lipid dye fluorescence (red, detected in 660/680 nm laser-filter unit), or DNA dye fluorescence (green, detected in 488/500 nm laser-filter unit) by NTA. A blue line is the fitted curve of the normal distribution to the experimental data. (B) The total concentration of particles detected by each signal with 50 ∼ 500 nm in size.

**Figure S5.**
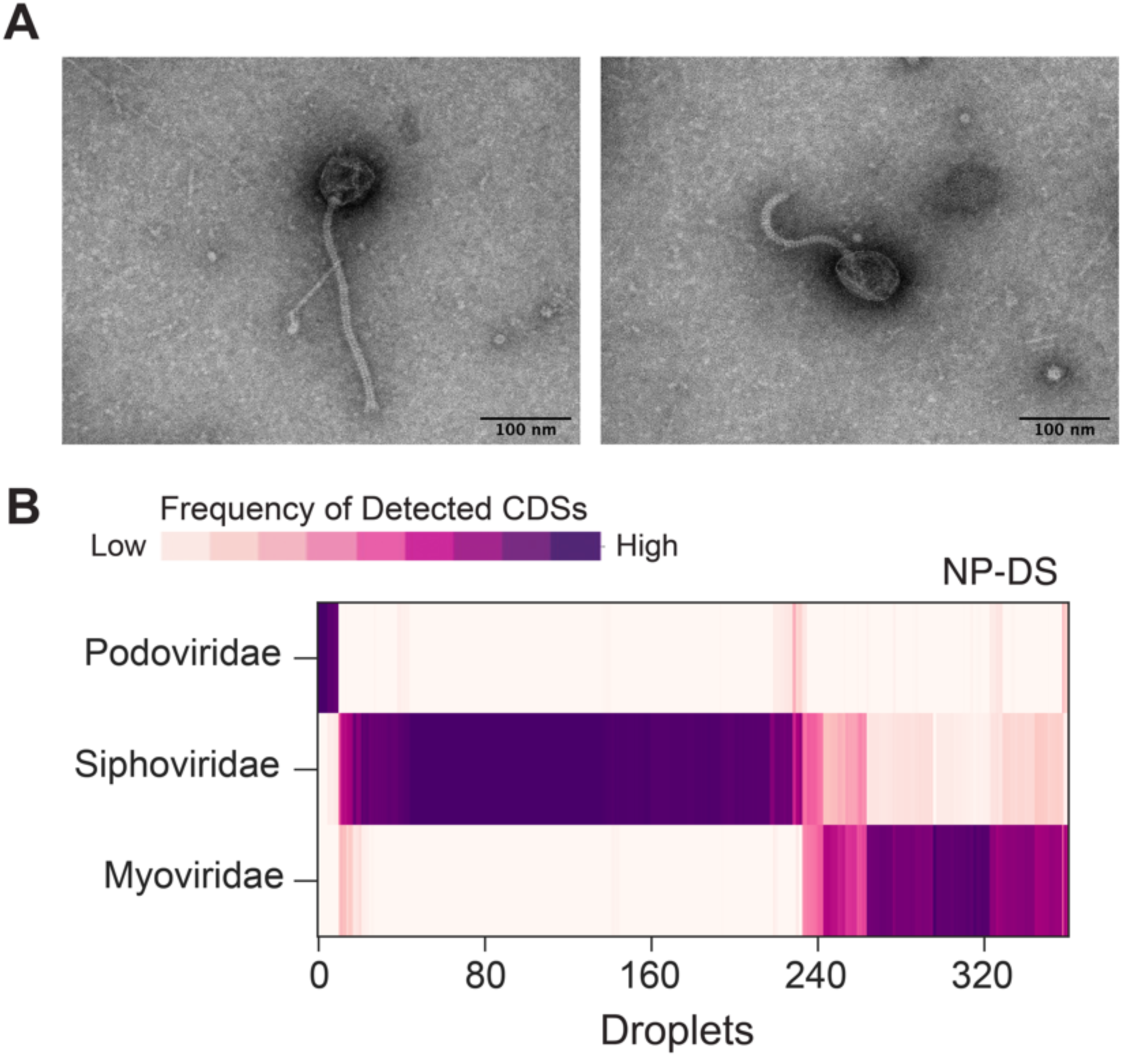
Phage particles in an isolated nanoparticle sample from the dental plaque. (A) The typical TEM micrographs of phage-like particles isolated from the dental plaque of periodontal patients. (B) Taxonomic profiles of phage-derived CDSs detected in NP-DS. Class-level taxonomic classification of viral CDSs (vCDSs) detected in NP-DS was shown as a heatmap. The color of the heatmaps showed the length of CDSs assigned to each taxonomy relative to the total length of vCDS detected in each droplet. We used NCBI taxonomy for classification. Phage species whose maximum frequency of detected CDS length in each droplet is less than 0.4 were not displayed.

**Figure S6.**
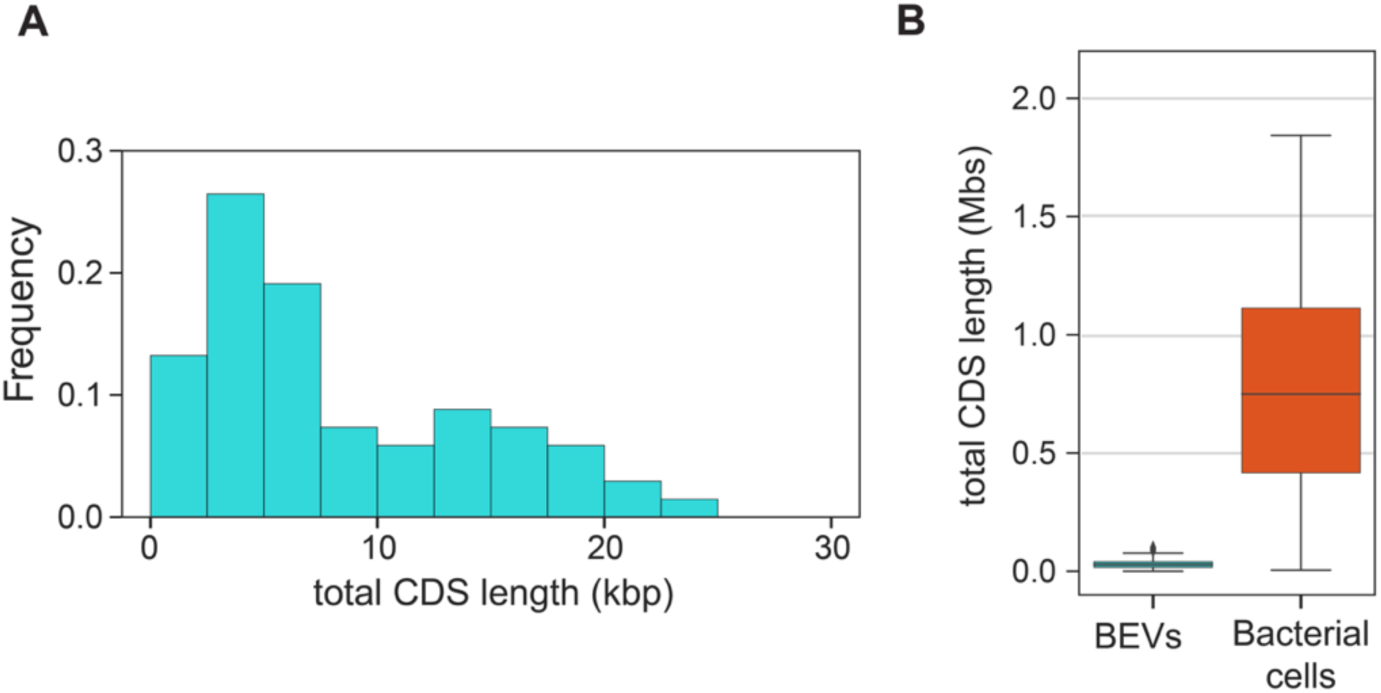
Total detected CDS length in BEV-containing droplets. (A) The distribution of total length of detected bCDS region in BEV-containing droplets. (B) Box plots of total bCDS length in BEV-containing droplets (NP-DS) and bacterial cells containing droplets (cell-DS). Medians and outliers were shown as gray lines and diamonds each.

**Figure S7.**
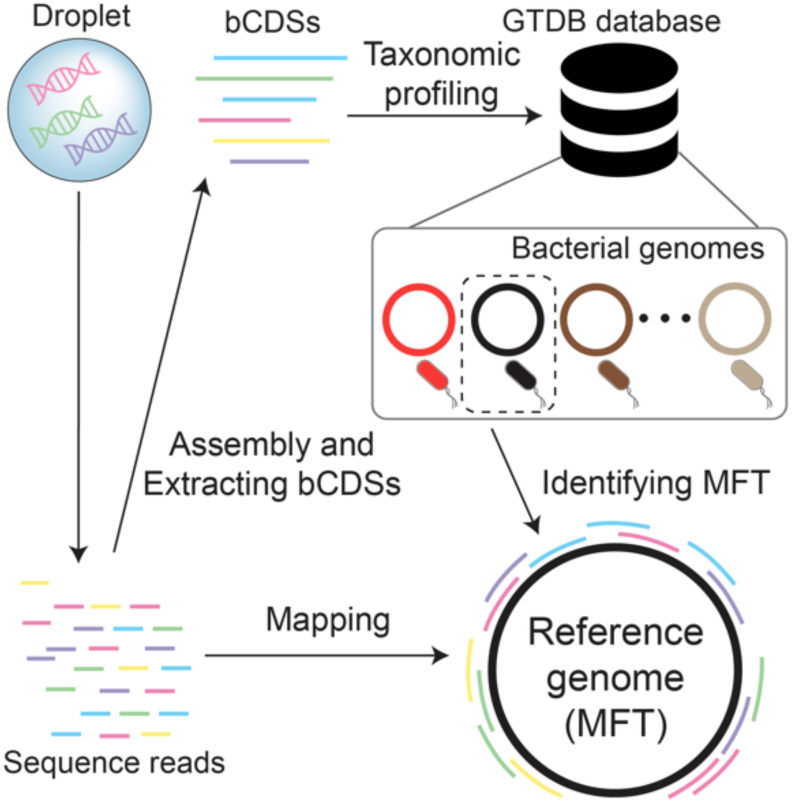
A schematic of the analysis for identifying MFT and mapping on the reference genome. For each droplet, the most frequently detected bacterial taxon (MFT) was identified in GTDB taxonomy based on the annotated information of bCDSs (for details, see Materials and Methods). Then the read sequences were mapped to the corresponding assembly genome of the MFT.

**Figure S8.**
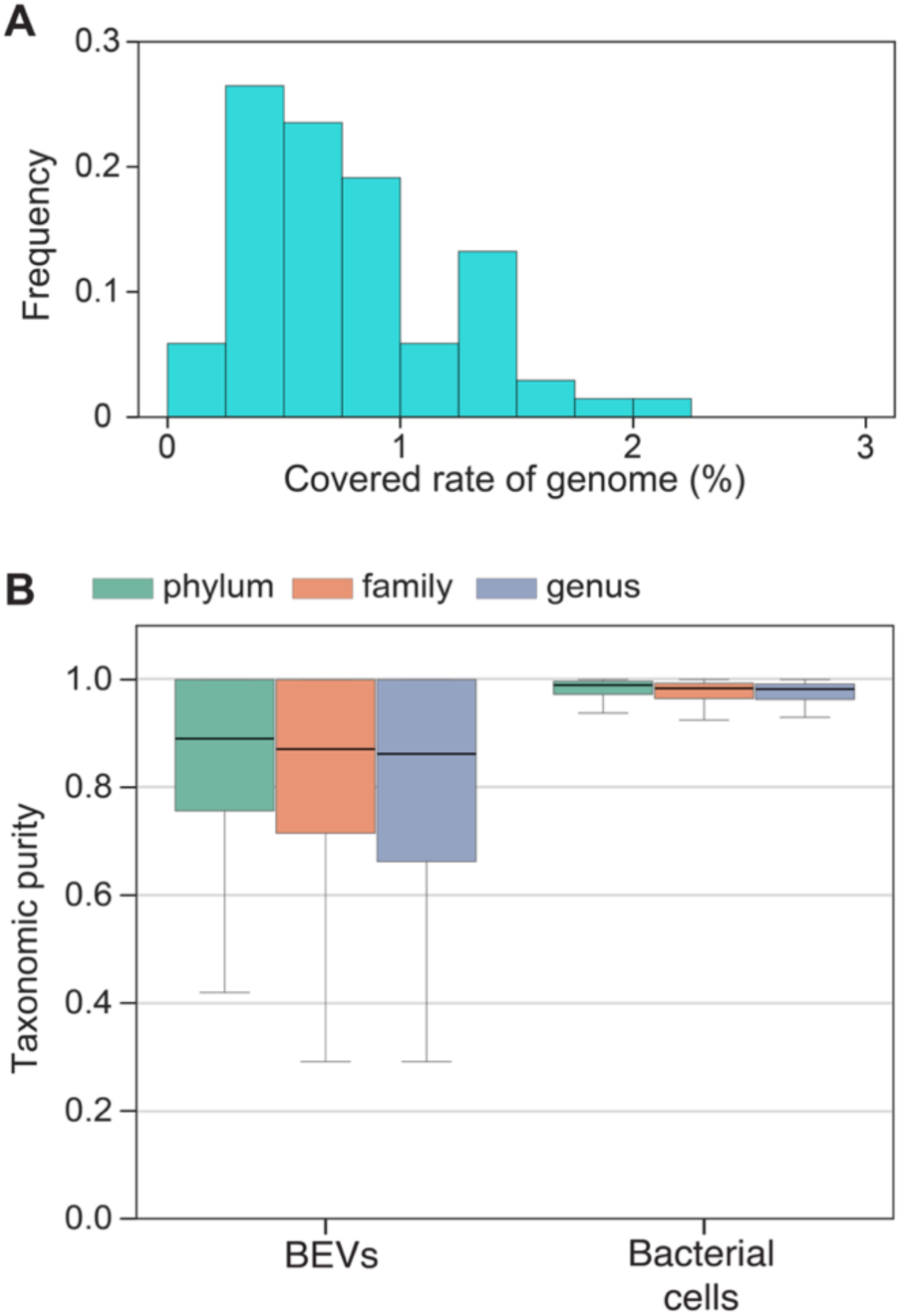
Mapping status of sequence reads in BEV-containing droplets. (A) The distribution of the covered rate of the genome of MFTs by sequence reads in each BEV-containing droplet. (B) Taxonomic purity of detected CDSs in BEVs isolated from the dental plaque. Here, the most frequently detected taxon (MFT) to which BEV-derived CDSs were assigned was first determined, and then the percentage of CDSs length attributed to the MFT in each droplet was calculated and plotted as a boxplot (we call this parameter “purity” of CDSs). We quantified this parameter for the GTDB taxonomic profile in phylum, class, and genus levels. Medians and outliers were shown as gray lines and diamonds each.

**Table S1.** Enriched CDSs in NP-DS of *P. gingivalis* BEVs. Enriched CDSs of *P. gingivalis* BEVs, which were screened by binominal test (p-value < 0.01). Those screened genes were identical to those screened by the threshold of 75 percentile + 2*IQR (interquartile range).

| **Protein id** | **description** | **start position (bp)** | **end position (bp)** |
| --- | --- | --- | --- |
| WP_013815260.1 | hypothetical protein | 2062054 | 2063016 |
| WP_005874812.1 | tetratricopeptide repeat protein | 2057308 | 2058195 |
| WP_005874932.1 | hypothetical protein | 2064777 | 2066783 |
| WP_230456087.1 | 2TM domain-containing protein | 2058359 | 2058541 |
| WP_005874833.1 | mechanosensitive ion channel | 2056208 | 2057206 |
| WP_230456086.1 | T9SS type A sorting domain-containing protein | 2058703 | 2059659 |
| WP_005874913.1 | hypothetical protein | 2066813 | 2067403 |
| WP_004583626.1 | DNA-binding protein | 2051440 | 2052066 |
| WP_005874827.1 | L-threonylcarbamoyladenylate synthase | 2052478 | 2053053 |
| WP_005874926.1 | CRISPR-associated endonuclease Cas2 | 2069305 | 2069595 |
| WP_005874927.1 | hypothetical protein | 2073529 | 2073972 |
| WP_005874822.1 | sugar transferase | 2053050 | 2054456 |
| WP_005874906.1 | type III-B CRISPR module RAMP protein Cmr6 | 2072771 | 2073532 |
| WP_005874912.1 | type III-B CRISPR module-associated protein Cmr3 | 2074684 | 2075874 |
| WP_010955945.1 | IS5 family transposase | 216542 | 217627 |
| WP_005874916.1 | CRISPR-associated endonuclease Cas1 | 2069615 | 2072710 |
| WP_010956463.1 | type III-B CRISPR module RAMP protein Cmr4 | 2073969 | 2074670 |
| WP_010955941.1 | IS4-like element IS1598 family transposase | 199227 | 200384 |
| WP_005873516.1 | threonine-phosphate decarboxylase CobD | 1239071 | 1240078 |
| WP_005873512.1 | cob(I)yrinic acid a,c-diamide adenosyltransferase | 1241563 | 1242129 |
| WP_005873514.1 | cobyrinate a,c-diamide synthase | 1242152 | 1243471 |
| WP_012458603.1 | DUF6261 family protein | 2060211 | 2061263 |
| WP_010956459.1 | DUF6261 family protein | 2063358 | 2064410 |
| WP_010956028.1 | IS5-like element ISPg8 family transposase | 502077 | 503162 |
| WP_005874030.1 | MULTISPECIES: histidinol phosphate phosphatase | 611806 | 612324 |
| WP_005874035.1 | purine nucleoside phosphorylase I, inosine and guanosine-specific | 615600 | 616421 |
| WP_005873715.1 | histidinol-phosphate aminotransferase | 623025 | 623531 |
| WP_005874530.1 | thiamine-phosphate kinase | 689154 | 690194 |
| WP_005873566.1 | adenosylcobinamide amidohydrolase | 696973 | 698136 |
| WP_004342769.1 | site-specific integrase | 878721 | 879923 |
| WP_004342772.1 | hypothetical protein | 879930 | 880292 |
| WP_014709482.1 | helix-turn-helix transcriptional regulator | 881003 | 881272 |
| WP_004584269.1 | ABC transporter permease | 1095538 | 1096284 |
| WP_005873460.1 | ATP-binding cassette domain-containing protein | 1096303 | 1097037 |
| WP_005873451.1 | T9SS C-terminal target domain-containing protein | 1097132 | 1098505 |
| WP_004584372.1 | S4 domain-containing protein | 1236875 | 1237327 |
| WP_010956235.1 | adenosylcobinamide-phosphate synthase CbiB | 1238047 | 1239057 |
| WP_005874841.1 | PH domain-containing protein | 1374934 | 1375389 |
| WP_004342775.1 | nuclear transport factor 2 family protein | 1534837 | 1535244 |

**Table S2.** Enriched gene categories in NP-DS of *P. gingivalis* BEVs. Enriched GO terms in the enriched CDSs in the BEVs were statistically screened (hypergeometric distribution test, p-value < 0.05). GO terms were grouped into three categories: M: Molecular functions; B: Biological processes; and C: Cellular components.

| GO id | category | name | Members<br>in original<br>genome | Members<br>in BEVs | p-value |
| --- | --- | --- | --- | --- | --- |
| GO:0009236 | P | cobalamin biosynthetic process | 13 | 4 | 5.02E-05 |
| GO:0048472 | F | threonine-phosphate decarboxylase activity | 2 | 2 | 0.000314 |
| GO:0043571 | P | maintenance of CRISPR repeat elements | 4 | 2 | 0.00185 |
| GO:0051607 | P | defense response to virus | 5 | 2 | 0.00304 |
| GO:0006313 | P | DNA transposition | 11 | 2 | 0.0157 |

**Table S3.** Enriched gene categories in bulk-BEV sequencing of *P. gingivalis*. Enriched GO terms screened in the bulk-BEV sequencing data (hypergeometric distribution test, p-value < 0.05).

| GO id | category | name | Members<br>in original<br>genome | Members<br>in BEVs | p-value |
| --- | --- | --- | --- | --- | --- |
| GO:0009236 | P | cobalamin biosynthetic process | 15 | 10 | 1.28E-06 |
| GO:0015420 | F | ABC-type vitamin B12 transporter activity | 4 | 4 | 0.000243 |
| GO:0043171 | P | peptide catabolic process | 6 | 4 | 0.00297 |
| GO:0032259 | P | methylation | 10 | 5 | 0.00442 |
| GO:0042026 | P | protein refolding | 4 | 3 | 0.00715 |
| GO:0042351 | P | 'de novo' GDP-L-fucose biosynthetic process | 2 | 2 | 0.0158 |
| GO:0048472 | F | threonine-phosphate decarboxylase activity | 2 | 2 | 0.0158 |
| GO:0000156 | F | phosphorelay response regulator activity | 2 | 2 | 0.0158 |
| GO:0070009 | F | serine-type aminopeptidase activity | 2 | 2 | 0.0158 |
| GO:0008641 | F | ubiquitin-like modifier activating enzyme activity | 2 | 2 | 0.0158 |
| GO:0022857 | F | transmembrane transporter activity | 13 | 5 | 0.0164 |
| GO:0006541 | P | glutamine metabolic process | 10 | 4 | 0.0276 |
| GO:0009399 | P | nitrogen fixation | 3 | 2 | 0.0435 |
| GO:0009408 | P | response to heat | 3 | 2 | 0.0435 |

**Table S4.** Enriched CDSs of *Alcaligenes faecalis* BEVs. CDSs detected in the enriched genomic region of *Alcaligenes faecalis* BEVs (p-value < 0.01, binominal test). CDS regions were extracted by Prokka and searched against the reference protein fasta file.

| **Protein id** | **description** | **start position (bp)** | **end position (bp)** |
| --- | --- | --- | --- |
| WP_042487686.1 | glycolate oxidase subunit GlcF | 981833 | 983053 |
| WP_042487683.1 | hypothetical protein | 983361 | 984668 |
| WP_009454582.1 | zeta toxin family protein | 1484000 | 1484545 |
| WP_009454580.1 | hypothetical protein | 1484541 | 1484726 |
| WP_042480022.1 | helix-turn-helix transcriptional regulator | 1484954 | 1485787 |
| WP_042480881.1 | efflux RND transporter periplasmic adaptor subunit | 1896312 | 1897547 |
| WP_042480884.1 | ABC transporter ATP-binding protein | 1897858 | 1898649 |
| WP_042487215.1 | 3-(3-hydroxy-phenyl)propionate transporter MhpT | 2690924 | 2692165 |
| WP_042487212.1 | hypothetical protein | 2692500 | 2693114 |
| WP_042486227.1 | chlorohydrolase family protein | 2981930 | 2983414 |
| WP_042486230.1 | tripartite tricarboxylate transporter substrate-binding protein | 2983444 | 2984421 |
| WP_042486802.1 | aspartate/glutamate racemase family protein | 3284430 | 3285188 |
| WP_042486928.1 | LysR substrate-binding domain-containing protein | 3285492 | 3286376 |
| WP_226791390.1 | ABC transporter permease | 3288210 | 3289034 |
| WP_042486807.1 | ABC transporter substrate-binding protein | 3289037 | 3290047 |
| WP_157766853.1 | acyltransferase family protein | 3290206 | 3292086 |
| WP_054513292.1 | Wzz/FepE/Etk N-terminal domain-containing protein | 3292644 | 3293687 |
| WP_042486816.1 | exopolysaccharide biosynthesis polyprenyl glycosylphosphotransferase | 3293751 | 3295019 |
| WP_042486818.1 | glycosyltransferase | 3295032 | 3295835 |
| WP_123794683.1 | hypothetical protein | 3295838 | 3296524 |
| WP_054513305.1 | IS5 family transposase | 3296656 | 3297621 |
| WP_054513289.1 | hypothetical protein | 3297683 | 3298210 |
| WP_042486050.1 | glycosyltransferase | 3298210 | 3299310 |
| WP_080723785.1 | N-acetyl sugar amidotransferase | 3299357 | 3300478 |
| WP_042486045.1 | AglZ/HisF2 family acetamidino modification protein | 3300478 | 3301236 |
| WP_042486042.1 | imidazole glycerol phosphate synthase subunit HisH | 3301236 | 3301850 |
| WP_052362964.1 | lipopolysaccharide biosynthesis protein | 3301850 | 3303094 |
| WP_042486039.1 | DegT/DnrJ/EryC1/StrS family aminotransferase | 3303137 | 3304228 |
| WP_042486036.1 | UDP-2-acetamido-3-amino-2,3-dideoxy-D-glucuronate N-acetyltransferase | 3304228 | 3304815 |
| WP_042486033.1 | Gfo/Idh/MocA family oxidoreductase | 3304815 | 3305759 |
| WP_080723784.1 | O-antigen ligase family protein | 3305990 | 3307249 |
| WP_042485889.1 | LysR substrate-binding domain-containing protein | 3379553 | 3380554 |
| WP_042485886.1 | tRNA 2-selenouridine(34) synthase MnmH | 3380547 | 3381638 |
| WP_042485881.1 | NosR/NirI family protein | 3381813 | 3384071 |
| WP_042485877.1 | TAT-dependent nitrous-oxide reductase | 3384137 | 3386038 |
| WP_226791393.1 | nitrous oxide reductase family maturation protein NosD | 3386049 | 3387386 |
| WP_042485874.1 | ABC transporter ATP-binding protein | 3387373 | 3388281 |
| WP_042485871.1 | ABC transporter permease | 3388288 | 3389115 |
| WP_042485868.1 | nitrous oxide reductase accessory protein NosL | 3389115 | 3389627 |
| WP_042485865.1 | FAD:protein FMN transferase | 3389663 | 3390667 |
| WP_042485861.1 | glucose-6-phosphate isomerase | 3390760 | 3392328 |
| WP_042485854.1 | LTA synthase family protein | 3393190 | 3394734 |
| WP_042485851.1 | SDR family NAD(P)-dependent oxidoreductase | 3394727 | 3395497 |
| WP_042485848.1 | glycosyltransferase | 3395487 | 3396956 |
| WP_042485845.1 | glycosyltransferase | 3396975 | 3398057 |
| WP_042485842.1 | glycosyltransferase family 4 protein | 3398057 | 3399370 |

**Table S5.** Enriched gene categories of *Alcaligenes faecalis* BEVs. Enriched GO terms in the enriched CDSs in the BEVs of *Alcaligenes faecalis* were statistically screened (hypergeometric distribution test, p-value < 0.05).

| GO id | category | name | Members in original genome | Members in BEVs | p-value |
| --- | --- | --- | --- | --- | --- |
| GO:0009243 | P | O antigen biosynthetic process | 9 | 5 | 9.30E-10 |
| GO:0000271 | P | polysaccharide biosynthetic process | 4 | 4 | 1.44E-09 |
| GO:0006065 | P | UDP-glucuronate biosynthetic process | 3 | 3 | 2.59E-07 |
| GO:0009103 | P | lipopolysaccharide biosynthetic process | 15 | 3 | 0.000112 |
| GO:0071555 | P | cell wall organization | 46 | 4 | 0.000198 |
| GO:0000107 | F | imidazoleglycerol-phosphate synthase activity | 4 | 2 | 0.000258 |
| GO:0000105 | P | histidine biosynthetic process | 14 | 2 | 0.00375 |
| GO:0016829 | F | lyase activity | 18 | 2 | 0.0062 |

**Dataset S1 (separate file .xlsx format).** Enriched CDSs in bulk BEV-sequencing of *P. gingivalis* BEVs. We screened genes whose FPKM were larger than the threshold (75 percentile + 2*IQR (interquartile range).

**Dataset S2 (separate file .xlsx format).** A list of all possible sources of contaminant DNA. All the species information in GTDB and NCBI formats were based on the listed taxa in either of Salter *et al.,* 2014 or Poore *et al.,* 2020 (14, 15). The DNA fragments that were taxonomically assigned to one of the listed accession numbers were eliminated.

